# Presynaptic Release Probability Determines the Need for Sleep

**DOI:** 10.1101/2025.10.16.682770

**Authors:** Yifan Wu, Keimpe Wierda, Katlijn Vints, Yu-Chun Huang, Valerie Uytterhoeven, Sahil Loomba, Fran Laenen, Marieke Hoekstra, Miranda C Dyson, Sheng Huang, Chengji Piao, Jiawen Chen, Sambashiva Banala, Chien-Chun Chen, El-Sayed Baz, Luke Lavis, Dion Dickman, Natalia V Gounko, Stephan Sigrist, Patrik Verstreken, Sha Liu

**Affiliations:** VIB-KU Leuven, Center for Brain & Disease Research, Leuven, 3000, Belgium; KU Leuven, Department of Neurosciences, Leuven Brain Institute, Leuven, 3000, Belgium; VIB-KU Leuven, Center for Brain & Disease Research, Electrophysiology Unit, Leuven, 3000, Belgium; VIB-KU Leuven, Bio Imaging Core, Leuven, 3000, Belgium; Max Planck Institute for Brain Research, Department of Connectomics, Frankfurt, 60438, Germany; Institute for Biology/Genetics, Freie Universität Berlin, Berlin, 14195, Germany; Department of Neurobiology, University of Southern California, Los Angeles, CA 90089, USA; Janelia Research Campus, Howard Hughes Medical Institute, Ashburn, VA 20174, USA; Suez Canal University, Zoology Department, Ismailia, 41522, Egypt

## Abstract

Sleep is universal among animals with synapses, yet the synaptic functions determining the need for sleep remain elusive. By directly measuring synaptic transmission at anatomically defined synapses in *Drosophila*, we found that synaptic strength remained stable or declined after sleep deprivation in a circuit-specific manner. In contrast, presynaptic release probability (P_r_) consistently decreased with sleep loss across circuits and species, stemming from reduced Ca^2+^ influx or weakened vesicle–channel coupling at presynaptic terminals, and recovered after sleep. Bidirectional manipulations of P_r_ altered sleep pressure, establishing a causal relationship between presynaptic function and sleep need. Non-synaptic sleep-regulatory signaling pathways consistently modulate P_r_ but not synaptic strength. Thus, our findings identify P_r_, rather than synaptic strength, as the conserved synaptic substrate underlying sleep need.

## Main Text

Sleep is an evolutionarily conserved behavior essential for maintaining optimal brain functions. Emerging evidence suggests that the restorative effects of sleep are tightly linked to its ability to adjust synaptic functions (*1–4*). The primary function of synapses is to transmit information between neurons through synaptic transmission, whereby presynaptic activity triggers neurotransmitter release and evokes postsynaptic responses. Yet it remains unclear which specific aspects of this process are consistently modulated across sleep and wake states.

The Synaptic Homeostasis Hypothesis (SHY) posits that wakefulness leads to a net increase in synaptic strength, while sleep globally downscales this heightened synaptic strength to maintain cellular and network stability (*1*). Previous attempts to evaluate SHY have relied primarily on structural and molecular observations, such as changes in synaptic morphology (*5–8*) and gene expression (*9–13*), which are commonly used as indirect proxies for synaptic strength. However, these associations have been established primarily under baseline conditions (*14*, *15*), without behavioral manipulations such as sleep deprivation, leaving their relevance to sleep and wakefulness uncertain. Moreover, although structural and molecular changes often accompany synaptic remodeling, they do not reliably reflect changes in synaptic strength (*16*, *17*). Direct functional measurements of synaptic strength across defined sleep–wake conditions are rare, and those that exist have often relied on spontaneous activity (*18*, *19*) rather than evoked responses from defined inputs—particularly at anatomically identified synapses—leaving the central functional predictions of SHY largely untested. Furthermore, even among the limited functional studies, most studies have focused on postsynaptic mechanisms (*10*, *12*), whereas the impact of sleep on presynaptic function remains poorly understood.

To address these gaps, we examined how sleep and wakefulness influence synaptic transmission in *Drosophila melanogaster* and mice. In flies, the stereotyped connectivity of defined circuits (*20*, *21*) allowed us to directly measure and compare synaptic strength at the same anatomically identified synapses across individuals. In both species, we further assessed presynaptic function by measuring release probability, a critical parameter indicating how effectively presynaptic activity triggers neurotransmitter release. Together, these experiments present a direct functional framework for examining how synaptic transmission, particularly presynaptic mechanisms, is modulated across sleep–wake cycles.

### Presynaptic release probability reflects sleep–wake history

To directly examine whether synaptic strength reflects sleep–wake history, as predicted by the Synaptic Homeostasis Hypothesis (SHY), we performed whole-cell patch-clamp recordings in the adult *Drosophila* brain following three defined states (Fig. 1, A and B): undisturbed sleep during the night (S), sleep deprivation induced by 10–12 hours of gentle, closed-loop mechanical stimulation during the dark phase (SD), and recovery sleep following deprivation (RS). We recorded from excitatory synapses in two brain regions that have been widely used to investigate synaptic physiology in *Drosophila* (*22–24*). In the antennal lobe, which processes olfactory information, we examined synaptic transmission from olfactory receptor neurons (ORNs) to projection neurons (PNs) in the VA1d glomerulus (Fig. 1A). In the mushroom body, the primary center for learning and memory, we recorded from synapses connecting Kenyon cells (KCs) to dorsal paired medial neurons (DPMs) (Fig. 1B).

**Fig. 1.**
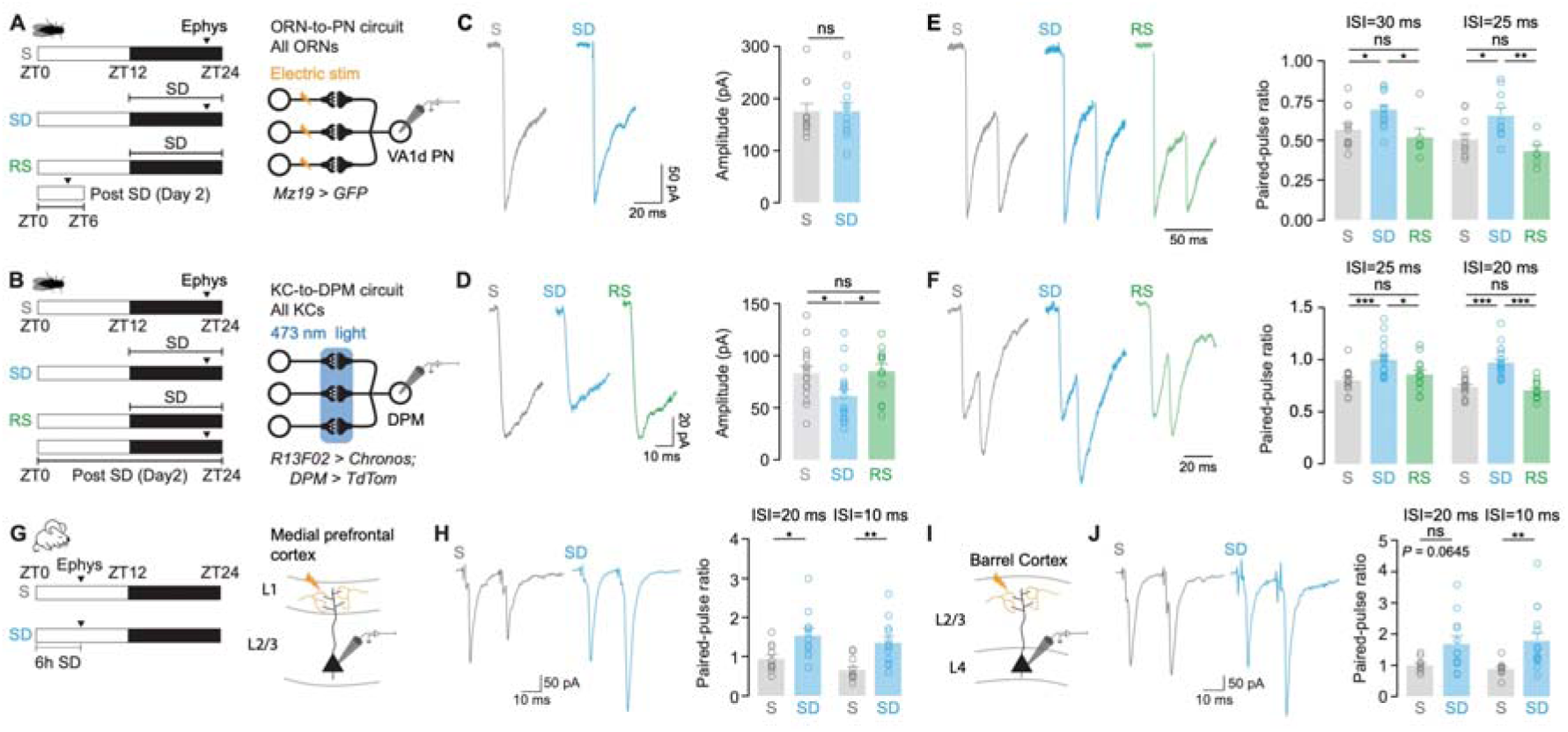
Sleep deprivation reduces presynaptic release probability and induces circuit-specific changes in synaptic strength and postsynaptic responses. (**A**) Flies were assigned to three groups: sleep (S), sleep-deprived (SD), and recovery sleep (RS). S and SD flies were recorded at ZT22 following undisturbed sleep or 10–12□h of sleep deprivation (ZT12–ZT24), respectively. RS flies were recorded at ZT4, following 12□h of SD on the previous day and 4–6□h of recovery sleep. Whole-cell recordings (black arrowhead) were performed from VA1d projection neurons (PNs) and lasted 2h. Axon bundle of olfactory receptor neurons (ORNs) were electrically stimulated; VA1d projection neurons (PNs) were labeled with GFP (*Mz19 > GFP*). (**B**) Whole-cell recordings (black arrowhead) were performed from dorsal paired medial (DPM) neurons, starting at ZT22 in S and SD flies, or ZT22 on the following day in RS flies. Kenyon cells (KCs) expressing Chronos (*R13F02*□*>*□*Chronos*) were optogenetically stimulated with 473 nm light; DPM neurons were labeled with tdTomato (*DPM > tdTom*). (**C**) Left: Representative excitatory postsynaptic currents (EPSCs) from S and SD groups, evoked by stimulating olfactory receptor neuron (ORN) afferents. Right: Mean EPSC amplitudes show no significant difference between groups (S: n = 12 from 12 flies; SD: n = 10 from 10 flies). (**D**) Left: Representative EPSCs from DPM neurons in S, SD, and RS groups following optogenetic stimulation of KCs. Right: Mean amplitudes are significantly reduced in SD and restored by RS (S: n = 16 from 16 flies; SD: n = 18 from 18 flies; RS: n = 13 from 13 flies). (**E**) Left: Representative normalized paired EPSCs from VA1d PNs in S, SD, and RS groups. Right: Paired-pulse ratio (PPR) is increased in SD and restored by RS. PPR was quantified at inter-stimulus intervals (ISIs) = 30 ms (S: n = 12 neurons from 12 flies; SD: n = 12 neurons from 12 flies; RS: n = 6 neurons from 6 flies) and 25 ms (S: n = 11 neurons from 11 flies; SD: n = 10 neurons from 10 flies; RS: n = 6 neurons from 6 flies). (**F**) Left: Representative normalized paired EPSCs from DPMs in S, SD, and RS groups. Right: PPR is increased in SD and restored by RS. PPR was quantified at ISIs = 25 ms (S: n = 16 neurons from 16 flies; SD: n = 18 neurons from 18 flies; RS: n = 13 neurons from 13 flies) and 20 ms (S: n = 17 neurons from 17 flies; SD: n = 18 neurons from 18 flies; RS: n = 17 neurons from 17 flies). (**G**) Experimental timeline and schematic of paired-pulse recording. Whole-cell recordings (black arrowhead) were performed from layer 2/3 pyramidal neurons in medial prefrontal cortex of sleep (S) and sleep-deprived (SD) mice. Layer 1 was electrically stimulated; SD mice were deprived of sleep for 6□h prior to recording. (**H**) Left: Representative traces of paired excitatory postsynaptic currents (EPSCs) recorded from S and SD mice. Right: Paired-pulse ratio (PPR) was increased in SD mice at inter-stimulus intervals (ISIs) of 20□ms (S: n = 11 neurons from 3 mice; SD: n = 11 neurons from 3 mice) and 10□ms (S: n = 12 neurons from 3 mice; SD: n = 12 neurons from 3 mice), indicating reduced presynaptic release probability. (**I**) Schematic of the barrel cortex recording configuration. Electrical stimulation was applied to layer 2/3, and EPSCs were recorded from layer 4 pyramidal neurons. The sleep deprivation timeline matched that used in (**G**). (**J**) Left: Representative traces of paired excitatory postsynaptic currents (EPSCs) recorded from S and SD mice. Right: PPR was increased in SD mice at a 10□ms interstimulus interval (S: n□=□10 neurons from 3 mice; SD: n□=□15 neurons from 3 mice), indicating reduced presynaptic release probability. At 20□ms, a similar trend was observed but did not reach statistical significance (*S*: *n*□=□10; *SD*: *n*□=□12 neurons from 3 mice; *P*□=□0.0645). Data are presented as mean□±□s.e.m. For statistical details, see Table S2. ns, not significant; *\*P*□*<*□*0.05, **P*□*<*□*0.01, ***P*□*<*□*0.001*.

Synaptic strength was assessed by measuring the amplitude of excitatory postsynaptic currents (EPSCs) evoked by presynaptic stimulation. At ORN-to-PN synapses in the antennal lobe, electrically evoked EPSC amplitudes remained stable across sleep and sleep deprivation (Fig. 1C), indicating that synaptic strength in this circuit is unaffected by sleep-wake states. In contrast, at KC-to-DPM synapses in the mushroom body, optogenetically evoked EPSC amplitudes using Chronos (*25*) were significantly reduced following SD and returned to baseline after 22 hours of RS (Fig. 1D). These results contradict SHY’s prediction of a general increase in synaptic strength with wakefulness. Instead, they reveal that synaptic strength is modulated in a circuit-specific manner across sleep–wake states.

Given the circuit-specific effects of sleep and wakefulness on synaptic strength, we asked whether another synaptic parameter might more consistently reflect sleep–wake state. We focused on presynaptic release probability (P_r_), which quantifies the likelihood that an incoming action potential triggers synaptic vesicle release, and is a key presynaptic determinant of synaptic strength. P_r_ was assessed using paired-pulse stimulation, a standard electrophysiological method (*26*) in which two closely spaced stimuli are delivered to the presynaptic terminal. The paired-pulse ratio (PPR), defined as the amplitude of the second EPSC relative to the first, is inversely related to P_r_ (*26*). In both ORN-to-PN and KC-to-DPM synapses, SD induced a robust and consistent increase in PPR across inter-stimulus intervals (ISIs; Fig. 1, E and F), indicating a reduction in P_r_. Notably, this reduction was fully reversed by recovery sleep, yet with circuit-specific timing: P_r_ returned to baseline within ∼6□hours at ORN-to-PN synapses but required approximately 22 □hours at KC-to-DPM synapses (Fig. 1, E and F). These observations demonstrate that, unlike synaptic strength, P_r_ shows consistent, reversible modulation across sleep–wake states in both circuits.

To determine whether sleep–wake–dependent changes in P_r_ also occur in mammals, we measured PPR in two excitatory pathways in the mouse cortex: layer 1 inputs onto layer 2/3 pyramidal neurons in the medial prefrontal cortex (mPFC, Fig. 1G) and layer 2/3 inputs onto layer 4 neurons in the barrel cortex (BC, Fig. 1I). These cortical regions have been used in prior studies examining sleep-dependent synaptic changes, including those supporting SHY (*5*, *27*). Consistent with our findings in flies, SD significantly increased PPR in both cortical pathways, indicating a decrease in P_r_. In mPFC, PPR was elevated at both 10-ms and 20-ms ISIs (Fig. 1H). In BC, PPR was significantly increased at 10-ms ISI, with a comparable but non-significant trend at 20 milliseconds (*P* = 0.0645, two-way repeated-measures ANOVA with Šidák test; Fig. 1J). Unlike the anatomically defined cholinergic excitatory circuits in *Drosophila*, these glutamatergic cortical pathways integrate heterogeneous excitatory inputs from multiple cortical sources. Yet, both species and circuit types exhibit parallel modulation of presynaptic function: wakefulness reduces P_r_ across evolutionarily divergent excitatory synapses. Such cross-species convergence points to presynaptic function as a key locus for sleep–wake modulation of synaptic transmission.

Previous studies have reported correlative changes in postsynaptic structure or receptor expression across sleep–wake states(*10*, *12*), but direct functional measurements remain limited. To directly assess postsynaptic function in *Drosophila*, we used photolysis-based uncaging of PANic2 (*28*), a caged nicotine analog, in the presence of tetrodotoxin (1□μM) to block presynaptic activity (fig. S1A). Wide-field illumination was used to uniformly activate all nicotinic acetylcholine receptors across the postsynaptic membrane (fig. S1, B and D). In VA1d PNs, uncaging-evoked currents were unaffected by SD (fig. S1C), whereas DPM neurons exhibited a significant reduction in postsynaptic response amplitude following SD (fig. S1E), aligning with the circuit-specific changes observed in synaptic strengths.

Collectively, our functional analyses reveal that P_r_, rather than synaptic strength, reliably encodes sleep–wake history across circuits and species. This differs from the SHY, which proposes global synaptic strength changes during sleep and wakefulness. Instead, presynaptic release probability stands out as the primary synaptic parameter modulated by sleep and wakefulness, and as a potential physiological substrate of sleep need at synapses.

### Sleep facilitates Ca^2+^-dependent synaptic vesicle release

Given that presynaptic release probability (P_r_) was consistently decreased across different synapses following sleep deprivation (SD), we next investigated the cellular mechanisms responsible for this reduction. Neurotransmitter release is a Ca²^+^-dependent process, whereby membrane depolarization at presynaptic terminals triggers Ca²□ influx through voltage-gated Ca^2+^ channels (VGCCs). This influx results in a localized elevation in Ca²□ concentration, driving synaptic vesicle (SV) exocytosis and subsequent neurotransmitter release in proximity to VGCCs. Thus, P_r_ is shaped by both the magnitude of evoked Ca^2+^ influx and the spatial coupling between VGCCs and SVs.

To assess whether reduced P_r_ after SD reflects altered Ca²□ influx, disrupted vesicle–Ca^2+^ channel coupling, or both, we first measured presynaptic Ca²□ influx at VA1d ORN terminals. Specifically, Syt1-GCaMP8m, a SV–tethered Ca^2+^ indicator, was co-expressed with the optogenetic actuator Chrimson (*25*) in Or88a-expressing ORNs, which project axons to the VA1d glomerulus. Presynaptic Ca^2+^ transients were then measured at their axon terminals in response to optogenetic stimulation (Fig. 2A). Surprisingly, Ca^2+^ responses evoked by 5-ms optogenetic activation were comparable between sleep and SD groups (Fig. 2, B and C), indicating that SD-induced reduction in P_r_ does not result from diminished presynaptic Ca²□ influx, at least at ORN terminals targeting the VA1d glomerulus.

**Fig. 2.**
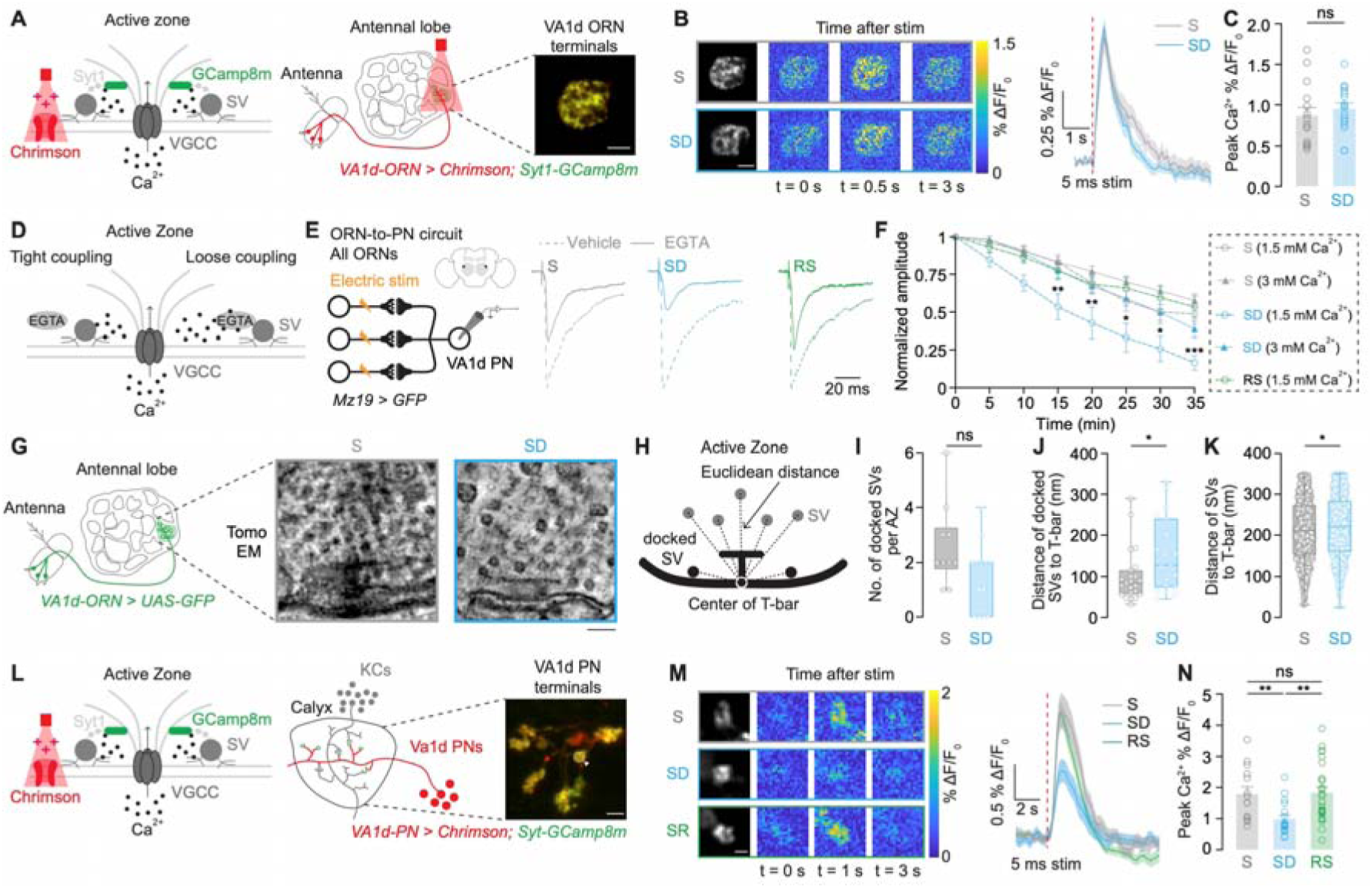
Sleep enhances presynaptic release probability via strengthened SV–VGCC coupling and increased Ca^2+^ influx. (**A**) All-optical approach for measuring presynaptic Ca^2+^ influx at VA1d olfactory receptor neuron (ORN) terminals. Left: at the presynaptic active zone, Chrimson-mediated membrane depolarization drives Ca^2+^ entry through voltage-gated Ca^2+^ channels (VGCCs), while Syt1-GCaMP8m, targeted to synaptic vesicle (SV) via Synaptotagmin-1 (Syt1), reports dynamic of local Ca^2+^ concentrations. Middle: anatomical diagram showing expression of Chrimson in VA1d ORNs projecting from the antenna to the antennal lobe. Ca^2+^ imaging was performed in the VA1d glomerulus, where presynaptic terminals reside. Right: Two-photon image showing Chrimson and Syt1-GCaMP8m fluorescence in VA1d ORN terminals within VA1d glomerulus. Scale bar: 10 μm. (**B**) Time-lapse two-photon images of GCaMP8m fluorescence in the axon terminals of VA1d ORNs in sleep (S) and sleep deprivation (SD) groups. Grayscale: baseline; pseudocolor: %ΔF/FL post-stimulation at 0, 0.5, 3 seconds(s). Scale bar: 10 μm. Right: Averaged Ca^2+^ responses across different animals in each group. (**C**) Peak Ca^2+^ transients in VA1d ORN terminals show no significant difference between S and SD groups. (S: n = 15 presynaptic terminals from 10 flies; SD: n = 16 presynaptic terminals from 11 flies). (**D**) Schematic of tight versus loose SV–VGCC coupling at the presynaptic active zone. In tight coupling (left), SVs are positioned close to VGCC, enabling rapid Ca^2+^-triggered release that is less sensitive to the slow Ca^2+^ chelator EGTA. In loose coupling (right), SVs are farther from VGCCs, allowing EGTA to more effectively buffer Ca^2+^ and reduce P_r_. (**E**) Left: Whole-cell recordings from VA1d projection neurons (PNs) during electrical ORN stimulation. Right: Representative traces of normalized EPSCs before (Vehicle control, DMSO; dashed line) and after application of 150□μM EGTA-AM (solid line) across S, SD, and recovery sleep (RS) groups under 1.5□mM extracellular Ca^2+^. (**F**) Time course of EGTA-AM effects on EPSCs at 1.5□mM (open circles) and 3□mM (solid triangles) extracellular Ca²□. Each Ca²□ condition was tested in a distinct set of flies (S: n = 9 at 1.5□mM, n = 9 at 3□mM; SD: n = 9 at 1.5□mM, n = 9 at 3□mM; RS: n = 10 at 1.5□mM only). SD flies showed greater EGTA sensitivity, particularly at 1.5 mM Ca²□, consistent with an increase in coupling distance between Ca²□ channels and SVs. This effect was reversed after 4–6□h of RS. Asterisks indicate significance between S and SD groups under 1.5□mM extracellular Ca²□. (**G**) Left: schematic showing expression of GFP in VA1d ORNs (*VA1d-ORN* > *UAS-GFP*) projecting from the antenna to the VA1d glomerulus. Right: Electron microscopy tomography (Tomo EM) of presynaptic boutons within the VA1d glomerulus from S and SD flies. Scale bar: 100 nm. (**H**) Schematic illustrating quantification of vesicle position relative to the center of the T-bar. (**I**) Number of docked vesicles per active zone (AZ) are reduced in SD compared to S, but the difference does not reach statistical significance (S: 10 T-bars, 2 flies; SD: 13 T-bars, 3 flies). (**J**) Distance from each docked SV to T-bar center is increased after SD (S: 23 vesicles; SD: 19 vesicles). (**K**) Distance from the T-bar center for all SVs within 350 nm is significantly increased in SD (S: 808 vesicles; SD: 1083 vesicles). (**L**) Left: Schematic of all-optical optogenetics and Ca^2+^ imaging in VA1d PN boutons. Right: Two-photon image showing Chrimson and Syt1-GCaMP8m fluorescence in VA1d PN boutons within the mushroom body calyx. A single bouton is marked by a dashed circle and arrowhead. Scale bar: 5 μm. (**M**) Time-lapse imaging of GCaMP8m fluorescence in a single bouton across conditions. Grayscale: baseline; pseudocolor: %ΔF/FL post-stimulation (0, 1, 3 s). Scale bar: 2 μm. Right: Averaged Ca^2+^ responses from individual boutons across different animals in each condition. (**N**) Peak Ca^2+^ transients in VA1d PN boutons decrease significantly after SD and return to baseline following 4-6 hours of RS (S: n = 14 from 5 flies; SD: n = 18 from 6 flies; RS: n = 26 from 5 flies). Data presented in panels (C), (F), and (N, left) are mean ± s.e.m. Box plots in panels (I), (J), and (K) display the median (center line), 25th and 75th percentiles (box), and minimum to maximum values (whiskers); Each data point represents a single neuron, synapse, or fly as indicated. For statistical details, see Table S2. ns, not significant; *\*P*□*<*□*0.05, **P*□*<*□*0.01, ***P*□*<*□*0.001*.

Since presynaptic Ca²□ influx at VA1d ORN terminals remained unchanged after SD, we next examined whether reduced P_r_ reflects altered spatial coupling between VGCCs and synaptic vesicles. To estimate coupling distance, we measured the reduction of evoked EPSCs by EGTA at VA1d ORN-to-PN synapses, a slow Ca²□ chelator that preferentially inhibits release from vesicles loosely coupled to Ca²□ channels and thus spatially more separated from VGCC(*29–31*) (Fig. 2D). Synaptic responses in sleep-deprived (SD) flies exhibited significantly greater sensitivity to EGTA than those in sleep (S) controls (Fig. 2, E and F) under 1.5 mM extracellular Ca²□, consistent with increased coupling distance after sleep loss. This effect was reversed after 4–6□h of recovery sleep (RS) (Fig. 2, E and F), as EGTA sensitivity in RS flies was no longer significantly different from S controls (S vs. RS, *P* > 0.05,

Two-way repeated-measures ANOVA with Šidák test). Additionally, elevating extracellular Ca²□ to 3 mM rescued EPSC amplitude in SD flies to control levels, as no significant difference was observed between SD and S groups at this concentration (Fig. 2F; S vs. SD, *P* > 0.05, Two-way repeated-measures ANOVA with Šidák test), consistent with functional decoupling that can be overcome by enhanced Ca²□ availability. To determine whether this functional decoupling corresponds to ultrastructural alterations at the presynaptic terminal, we employed correlative light and electron microscopy (CLEM) combined with electron tomography to examine the ultrastructure of presynaptic boutons within the VA1d glomerulus at 1.1□nm isotropic resolution, focusing on synaptic vesicle organization (Fig. 2G). In *Drosophila* synapses, VGCCs cluster at the center of the active zone(*32*), beneath the T-bar, an electron-dense presynaptic structure that organizes synaptic vesicles (SV) and defines the site of neurotransmitter release. The spatial distribution of vesicles relative to the T-bar therefore serves as a structural correlate of SV–VGCC coupling (Fig. 2H). Although the number of docked vesicles per active zone showed a modest, insignificant reduction following SD (Fig. 2I; *P* = 0.081, Linear fixed model), we observed a significant shift in their distribution, with docked vesicles positioned farther from the T-bar center in SD compared to controls (Fig. 2J). A similar redistribution was observed across the entire vesicle population within 350 nm of the active zone (Fig. 2K). These structural rearrangements likely underlie the observed functional decoupling between SVs and VGCCs. Collectively, these functional and ultrastructural findings suggest that SD weakens the tight SV-VGCC coupling at ORN-to-PN synapses, thereby reducing P_r_ without affecting Ca²□ influx itself.

We next asked whether similar Ca²□-dependent mechanisms underlie P_r_ changes in other neuron types. To address this, we focused on VA1d projection neurons (PNs) (Fig. 2L), whose presynaptic boutons in the mushroom body calyx are sparsely distributed and large enough to permit single-bouton Ca^2+^ imaging, using the same strategy as in Fig. 2A. In contrast to ORN terminals, PN boutons exhibited a significant reduction in optogenetically evoked Ca^2+^ influx following SD, which fully recovered after 4–6 hours of recovery sleep (Fig. 2, M and N). This reduction in Ca^2+^ entry is unlikely to stem from altered neuronal excitability, as the intrinsic electrophysiological properties of VA1d PNs remained unchanged after SD (fig. S2). Because presynaptic Ca^2+^ influx directly determines P_r_, these findings also suggest that P_r_ is diminished in VA1d PN boutons after SD. Due to the sparse and stochastic connectivity between PNs and KCs, reliably targeting postsynaptic KCs for electrophysiological recordings is challenging, precluding direct assessment of SV–VGCC coupling in this circuit. Nevertheless, our results suggest that sleep loss disrupts Ca^2+^-dependent vesicle release in both ORN-to-PN and KC-to-DPM synapses, either by loosening SV-VGCC coupling or by impairing Ca^2+^ influx through VGCC, leading to a common functional outcome: reduced P_r_.

Considering the central role of presynaptic Ca^2+^ signalling in regulating P_r_, we investigated whether genetically impairing evoked Ca^2+^ entry at presynaptic terminals directly modulates sleep. Since the null mutant of *Cacophony* (*Cac*), the VGCC at the active zones, is lethal, we utilized a viable hypomorphic mutant (*33*), *Cac^exon7^*^Δ^, which has a deletion in exon 7. Paired-pulse analysis at neuromuscular junction (NMJ) of third-instar larvae revealed that *Cac^exon7^*^Δ^ mutants exhibited a significant increase in the PPR at 10- and 20-ms inter-stimulus intervals (ISI) (fig. S3A), indicating a reduction in P_r_. Given that SD consistently reduces P_r_ across circuits, we predicted that *Cac^exon7^*^Δ^ mutant, with constitutively lowered P_r_, would phenocopy a state of elevated sleep need. Indeed, *Cac^exon7^*^Δ^ flies showed a significant increase in daily sleep amount (fig. S3B), particularly during the night, accompanied by longer sleep bouts without a corresponding change in bout number (fig. S3B), suggesting consolidated sleep. As shown in fig. S4, *Cac^exon7Δ^* mutants were less aroused by mild mechanical stimulation compared to wild-type (WT) controls, while both genotypes responded robustly to strong stimulation, indicating that *Cac^exon7Δ^* mutants have an increased arousal threshold. These findings demonstrate that reduced presynaptic Ca^2+^ influx lowers P_r_ and elevates sleep drive, identifying presynaptic Ca^2+^ signaling as a critical cellular mechanism for encoding sleep need.

### Presynaptic release probability encodes sleep need

To determine whether the sleep-promoting effect observed in the *Cac^exon7^*^Δ^ mutant arises specifically from reduced presynaptic Cac levels or more generally from decreased presynaptic release probability (P_r_), we examined additional mutants targeting core components of the presynaptic release machinery known to affect P_r_. Specifically, we assessed sleep in null mutants for *Fife* (*34*), encoding a scaffolding protein essential for synaptic vesicle docking, and *Rim* (*35*, *36*), encoding a protein involved in vesicle priming and Ca^2+^ channel tethering at active zone (Fig. 3A). Both *Fife^AC^* and *Rim^Ex73^* mutants have been previously shown to exhibit reduced P_r_ in *Drosophila* (*37*). Consistent with this, we found that both null mutant lines exhibited significantly increased daily sleep amount compared to wild-type controls (Fig. 3B). This increase in sleep is not due to impaired locomotion, as waking activity (locomotor speed during wakefulness) was not decreased in either mutant (fig. S5A).

**Fig. 3.**
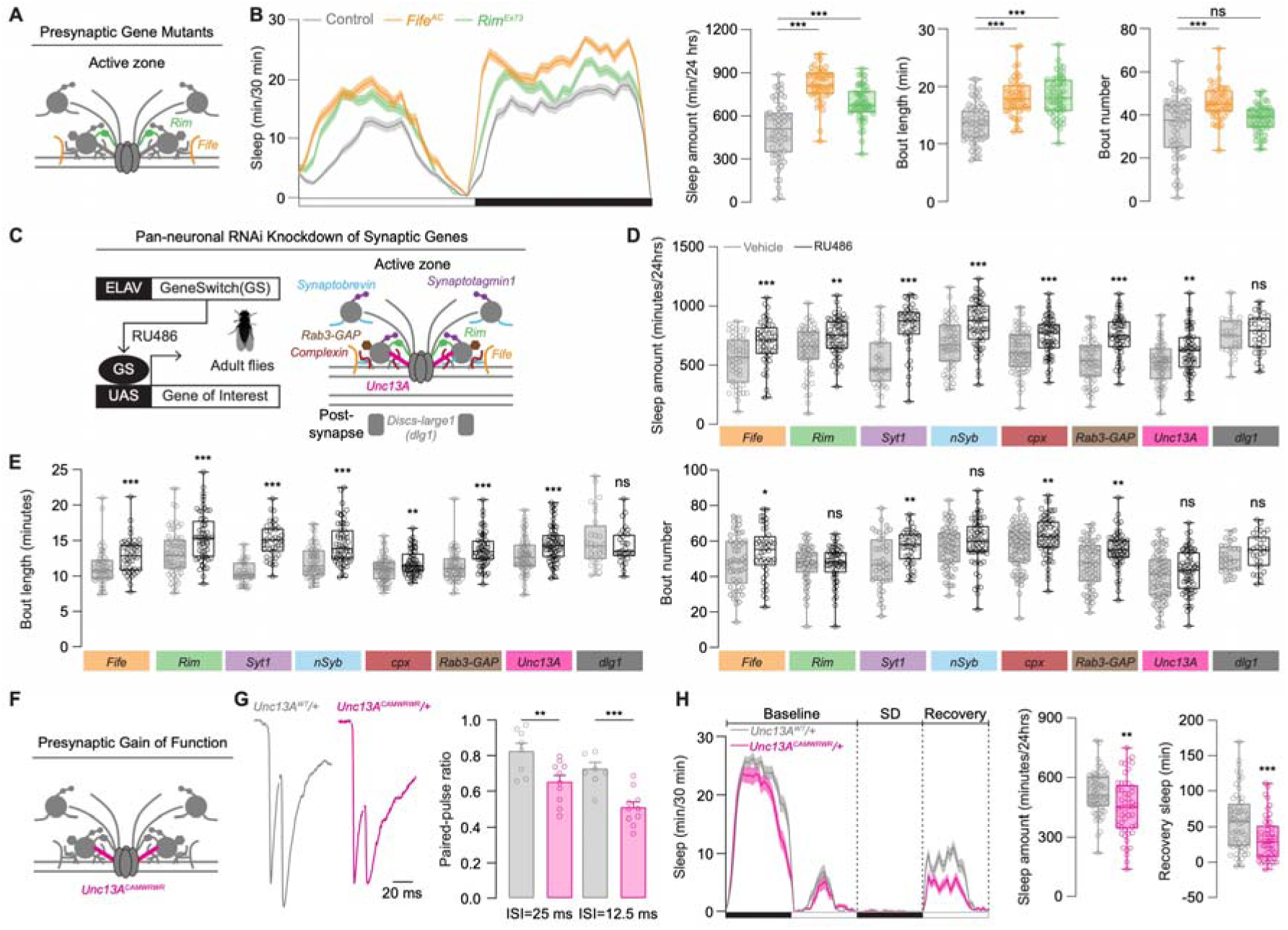
Manipulation of presynaptic release machinery bidirectionally regulates sleep. (**A**) Schematic of the presynaptic active zone highlighting key components of the presynaptic release machinery, including the proteins Fife and Rim, which regulate synaptic vesicle fusion and thus neurotransmitter release. Null mutants for each were used to assess their role in sleep regulation. (**B**) Sleep profiles and architecture. Left: Sleep profiles of *Fife^AC^*, *Rim^Ex73^*, and control flies, plotted in 30-min bins. Right: Daily sleep amount (min/24□hrs), mean sleep bout length (min), and number of sleep bouts. Both *Fife^AC^* and *Rim^Ex73^* mutants show significantly increased sleep and longer, more consolidated bouts. *Fife^AC^* (Orange, n=43 flies), *Rim^Ex73^* (Green, n=53 flies), and control (Gray, n=60 flies). (**C-E**), Pan-neuronal RNAi knockdown of presynaptic release machinery genes in adult flies. (**C**) Left: Schematic of the GeneSwitch (GS) system used to induce pan-neuronal RNAi knockdown in adult flies where *ELAV*-GS drives expression of *UAS-RNAi* constructs in neurons only after RU486 administration, enabling temporal control of gene silencing. Right: diagram of the synapse showing genes targeted by RNAi. Most genes (e.g., *Fife*, *Rim*, *Syt1*, *nSyb*, *cpx*, *Rab3-GAP*, *Unc13A*) act at the presynaptic active zone to regulate neurotransmitter release, while *dlg1* functions postsynaptically. (**D**) Total sleep amount (min/24□h) in flies fed with vehicle control or RU486. Knockdown of presynaptic release genes significantly increased sleep compared to controls, whereas knockdown of the postsynaptic gene *dlg1* had no effect. (**E**) Sleep bout length (left) and number of sleep bouts (right) following RU486 treatment. RNAi of presynaptic release genes led to longer, more consolidated sleep bouts. Sample sizes (vehicle, RU486): *Syt1* (n□=□38, 42 flies), *nSyb* (n□=□60, 57 flies), *cpx* (n□=□75, 64 flies), *Rab3-GAP* (n□=□60, 59 flies), *Unc13A* (n□=□69, 68 flies), *Fife* (n□=□50, 44 flies), *Rim* (n□=□59, 60 flies), *dlg1* (n□=□28, 30 flies). (**F**) Schematic illustrating the presynaptic active zone highlighting *Unc13A^CAMWRWR^*, a gain-of-function mutant enhancing presynaptic release probability. (**G**) Left: Representative normalized paired EPSCs from *Unc13A^CAMWRWR^/+* and *Unc13^WT^/+*. Right: Paired-pulse ratio is reduced in *Unc13A^CAMWRWR^/+* at 25 ms and 12.5 ms inter-stimulus intervals (ISI) under 0.5□mM extracellular Ca²□, indicating increased presynaptic release probability (*Unc13A^CAMWRWR^/+*: n = 8 neuron from 4 flies and *Unc13A^WT^/+*: n = 10 neurons from 6 flies). (**H**) Sleep profiles binned in 30-min intervals (left), total sleep duration over 24□hours (middle), and sleep recovered during the first 4□hours after 12□hours of mechanical sleep deprivation (right) for *Unc13A^CAMWRWR^/+* (n = 49) and *Unc13^WT^/+* (n = 60) flies. Data presented in panels (B, left), (G) and (H, left) are mean ± s.e.m. Box plots in panels (B), (D), (E), and (H) display the median (center line), 25th and 75th percentiles (box), and minimum to maximum values (whiskers); Each data point represents a single neuron or fly as indicated. For statistical details, see Table S2. ns, not significant; *\*P*□*<*□*0.05, **P*□*<*□*0.01, ***P*□*<*□*0.001*.

In addition, both lines showed prolonged sleep bouts, and *Fife^AC^*mutants also exhibited an increased number of sleep bouts (Fig. 3B). Together, these results suggest that a reduction in P_r_, regardless of the specific molecular mechanism, is sufficient to promote sleep.

To rule out the possibility that the observed sleep phenotypes resulted from impaired motor function, we performed cholinergic neuron-specific knockdowns of *Fife* and *Rim* using the *ChAT-T2A-GAL4* driver(*38*), which targets all cholinergic neurons while excluding motor neurons in *Drosophila* (fig. S6A). As fly motor neurons are glutamatergic and central excitatory neurons are predominantly cholinergic, this allowed us to assess excitatory circuit function without confounding motor output. These cholinergic neuron-specific knockdowns recapitulated the increased sleep duration and depth observed in the respective null mutants. Specifically, the daily sleep amount and sleep bout length were significantly increased compared to both Gal4 and UAS controls (fig. S6B), while waking activity was not reduced (fig. S5B), suggesting that the sleep phenotypes reflect genuine alterations in sleep need rather than motor impairments.

Next, we employed the RU486-inducible GeneSwitch system (*39*) to achieve adult-specific, pan-neuronal knockdown of a wide range of presynaptic genes (Fig. 3C). This approach bypassed developmental lethality and enabled temporal control over P_r_ disruption. As expected, adult-specific, pan-neuronal knockdown of *Fife* and *Rim* phenocopied the enhanced sleep phenotypes observed in their respective mutants and cholinergic knockdowns (Fig. 3, D and E). We next examined additional components of the presynaptic release machinery, including the synaptic vesicle fusion proteins *Synaptotagmin1 (Syt1)* (*40*, *41*)*, neuronal Synaptobrevin (nSyb)* (*42*)*, complexin (cpx)* (*43*, *44*)*, and Rab3-GAP* (*45*), as well as the active zone protein *Unc13A* (*46–48*). Knockdown of each of these genes resulted in a significant increase in total sleep amount compared to vehicle-fed controls (Fig. 3D). Sleep bout length increased across all genotypes, while *Syt1*, *cpx*, and *Rab3-GAP* knockdowns also exhibited a significant increase in bout number (Fig. 3E), reflecting a shift toward more consolidated sleep. Waking activity remained unchanged in all knockdown lines (fig. S5C), indicating that reduced P_r_ does not impair locomotion. In contrast, knockdown of *dlg1* (*49*), encoding a postsynaptic scaffolding protein, did not affect sleep (Fig. 3, D and E), indicating that the sleep-promoting effects observed are specific to presynaptic P_r_ disruption. Collectively, these findings demonstrate that reduced P_r_, achieved through various genetic manipulations of presynaptic components, is sufficient to elevate sleep need in *Drosophila*, highlighting the critical role of presynaptic function in sleep regulation.

A defining feature of sleep is the compensatory increase in sleep following SD, commonly termed rebound sleep or recovery sleep (RS). If P_r_ encodes sleep need, then constitutively elevating P_r_ should attenuate RS. To assess this, we examined a gain-of-function allele, *Unc13A^CaMWRWR^* (Fig. 3F), previously shown to exhibit elevated P_r_ in the neuromuscular junction (NMJ) (*50*). To determine whether this effect extends to central synapses, we evaluated P_r_ in the ORN-to-PN circuit in the adult brain. We found that flies carry one copy of *Unc13A^CaMWRWR^* displayed increased P_r_ under low extracellular Ca^2+^ conditions (0.5 mM), reflected by reduced PPR (Fig. 3G). Behaviourally, these flies exhibit significant reduction in baseline sleep and a markedly attenuated RS following 12 hours of SD (Fig. 3H), supporting the hypothesis that elevated P_r_ attenuates the accumulation of homeostatic sleep need.

Together, these findings support a model in which P_r_ at excitatory synapses serves as both a readout of accumulated and a determinant of sleep need. Reductions in P_r_, induced by either genetic perturbations or sleep deprivation, lead to an increased need for sleep, whereas elevations in P_r_ diminishes the homeostatic response to sleep loss. These results thus identify P_r_ as a key mechanistic substrate through which the nervous system encodes and determines sleep need.

### Distinct sleep-regulatory signaling pathways converge on P_r_

Multiple signaling pathways that regulate sleep and wakefulness have been identified, yet their direct effects on synaptic function remain unclear. Thus, we investigated whether such established regulators of sleep modulate synaptic functions, especially presynaptic release probability (P_r_) or synaptic strength, as part of their mechanism in sleep regulation.

Salt-Inducible Kinase 3 (*Sik3*), an evolutionarily conserved serine/threonine kinase, has been implicated in sleep regulation across species (*51*). A phosphorylation-deficient mutant form, *Sik3^S563A^* in flies and *Sik3^Sly^* in mice, promotes sleep (*51*). Using the RU486-inducible GeneSwitch driver, *ELAV-GS*, we pan-neuronally overexpressed *Sik3^S563A^* specifically in adult flies and observed increased sleep duration (Fig. 4, A and B) and bout length (fig. S7A), consistent with previous reports (*51*). Given *Sik3^Sly^* mice display altered phosphorylation states in multiple presynaptic proteins (*52*), we tested whether *Sik3^S563A^* overexpression modulates presynaptic function in flies. At ORN-to-PN synapses in the VA1d glomerulus, evoked EPSC amplitudes were unchanged by *Sik3^S563A^* overexpression (Fig. 4C), indicating stable synaptic strength. However, paired□pulse recordings showed a significant increase in paired□pulse ratio (PPR) at both 20□ms and 25□ms inter□stimulus intervals (Fig. 4D), indicating a reduction in P_r_. This selective modulation suggests Sik3 promotes sleep, at least in part, by downregulating P_r_, rather than increasing synaptic strength.

**Fig. 4.**
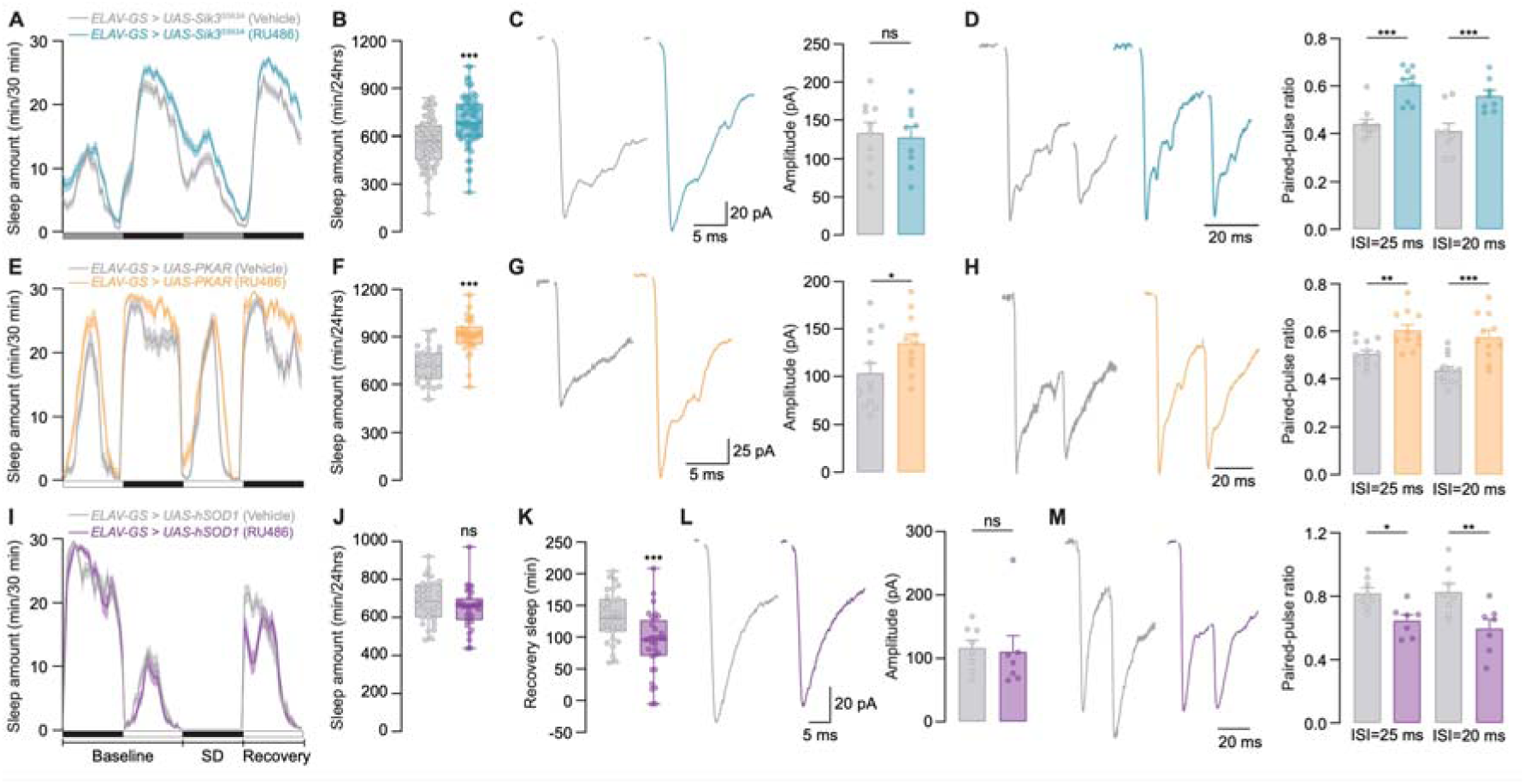
Sleep-regulatory signaling pathways modulate presynaptic release probability. (**A-D**) *Sik3^S563A^* was pan-neuronally overexpressed in adult flies using the RU486-inducible *ELAV-GeneSwitch (GS)* system to test whether its sleep-promoting effect involves changes in synaptic strength and presynaptic release probability (P_r_). (**A**) Sleep profiles of *ELAV-GS > UAS-Sik3^S563A^* flies in constant darkness, treated with vehicle or RU486, plotted in 30-min bins. (**B**) RU486-fed flies showed increased total sleep amount (min/24□hrs) and longer sleep bouts (min), with no change in bout number. Vehicle: n = 70□flies; RU486: n = 70 flies. (**C**) Representative evoked EPSC traces (Left) and mean EPSC amplitudes (Right) show no significant difference between groups (Vehicle: n = 10 neurons from 4 flies; RU486: n = 9 neurons from 3 flies) under 1.5□mM extracellular Ca^2+^. (**D**) Paired-pulse ratio (PPR) was significantly increased at 25□ms (Vehicle: n = 10 neurons from 4 flies; RU486: n = 9 neurons from 3 flies) and 20□ms (Vehicle: n = 9 neurons from 4 flies; RU486: n = 8 neurons from 3 flies) inter-stimulus intervals (ISIs) in RU486-fed flies under 1.5□mM extracellular Ca^2+^, indicating reduced P_r_. (**E–H**) *PKAR,* a regulatory subunit of protein kinase A (PKA), was pan-neuronally overexpressed in adult flies using the *GS* system to test whether its sleep-promoting effects involve changes in synaptic strength and P_r_. (**E**) Sleep profiles of *ELAV-GS > UAS-PKAR* flies, treated with vehicle or RU486, plotted in 30-min bins. (**F**) RU486-fed flies showed increased total sleep amount (min/24□hrs) and longer sleep bouts (min), with no change in bout number. Vehicle: n=29 flies; RU486: n=30□flies. (**G**) Representative evoked EPSC traces (Left) and mean EPSC amplitudes (Right) are significantly increased in RU486-fed flies (Vehicle: n=13 neurons from 4 flies; RU486: n = 11 neurons from 4 flies) under 1.5□mM extracellular Ca^2+^. (**H**) PPR was significantly increased at 25□ms and 20□ms ISIs in RU486-fed flies under 1.5□mM extracellular Ca^2+^ (Vehicle: n=13 neurons from 4 flies□; RU486: n = 11 neurons from 4 flies), indicating reduced P_r_. (**I-M**) Human *SOD1* (*hSOD1*) was pan-neuronally overexpressed in adult flies using the *GS* system to test whether its effect on sleep homeostasis is mediated through P_r_. (**I**) Sleep profiles of *ELAV-GS > UAS-hSOD1* flies, treated with vehicle or RU486, plotted in 30-min bins. (**J**) RU486-fed flies showed no change in baseline sleep amount. (**K**) Recovery sleep following sleep deprivation was significantly reduced in RU486-fed flies. (Vehicle: n = 31; RU486: n = 32 flies). (**L**) Representative evoked EPSC traces (Left) and mean EPSC amplitudes (Right) show no significant difference between groups (Vehicle: n=□8 neurons from 4 flies; RU486: n = 7 neurons from 4 flies) under 0.5□mM extracellular Ca^2+^. (**M**) PPR was significantly decreased at 25 ms and 20□ms ISIs in RU486-fed flies under 0.5□mM extracellular Ca^2+^ (Vehicle: n=□8 neurons from 4 flies; RU486: n = 7 neurons from 4 flies), indicating enhanced P_r_. Data presented in panels (A), (C), (D), (E), (G), (H), (I), (L) and (M) are mean ± s.e.m. Box plots in panels (B), (F), (J), and (K) display the median (center line), 25th and 75th percentiles (box), and minimum to maximum values (whiskers); Each data point represents a single neuron or fly as indicated. For statistical details, see Table S2. ns, not significant; *\*P*□*<*□*0.05, **P*□*<*□*0.01, ***P*□*<*□*0.001*.

Next, we examined protein kinase A (PKA), a conserved wake-promoting kinase. When we pan-neuronally overexpressed a regulatory subunit of the PKA regulatory subunit (PKAR) (*53*) specifically in the adult stage, thereby inhibiting PKA signalling, flies exhibited increased sleep duration (Fig. 4, E and F) and bout length (fig. S7B), as expected. In contrast to *Sik3^S563A^* overexpression, PKA inhibition significantly increased EPSC amplitude at ORN-to-PN synapses (Fig. 4G), indicating enhanced synaptic strength. Consistent with *Sik3^S563A^* overexpression, paired-pulse analysis in PKAR-overexpressing flies revealed reduced P_r_ (Fig. 4H). These results suggest that PKA promotes wakefulness, at least partly, through enhancing P_r_.

In addition to kinase-based signalling pathways such as Sik3 and PKA, metabolic state— particularly redox balance—has been implicated in sleep regulation (*54–57*). An influential hypothesis posits that sleep serves to counteract oxidative stress in the brain (*58*). Reactive oxygen species (ROS), which accumulate during wakefulness as metabolic by-products, can damage cellular components if not effectively neutralized. Sleep is thought to restore redox homeostasis by promoting antioxidant activity and preserving neural integrity (*58*). Although overexpression of antioxidant genes pan-neuronally has been shown to reduce baseline sleep in flies (*54*), its impact on the accumulation of sleep need remains unclear. To test this, we overexpressed human superoxide dismutase 1 (hSOD1) in adult neurons to reduce the ROS levels (*59*) and assessed recovery sleep (RS) following 12 hours of sleep deprivation (Fig. 4I). While baseline sleep was unaffected (Fig. 4J), hSOD1 overexpressing flies exhibited a marked reduction in RS (Fig. 4K), indicating decreased homeostatic sleep pressure. We next examined whether this effect correlates with changes in synaptic strength and/or P_r,_. EPSC amplitudes in hSOD1-overexpressing flies remained unchanged compared to controls (Fig. 4L), indicating stable synaptic strength at the ORN-to-PN synapse. In contrast, PPR was significantly reduced at 20- and 25-ms intervals (Fig. 4M), reflecting an increase in P_r_. These results indicate that enhanced oxidative stress clearance is associated with elevated P_r_, suggesting increased P_r_, rather than reduced synaptic strength, is the primary synaptic consequence of altered redox state, that potentially attenuating the accumulation of sleep need.

Taken together, these findings indicate that distinct sleep-regulatory pathways, including the sleep-promoting kinase SIK3, the wake-promoting kinase PKA, and neuronal redox state, converge on modulation of P_r_. In contrast, changes in synaptic strength are inconsistent across sleep-regulatory pathways, suggesting that synaptic strength plays a comparatively minor role in sleep regulation. Thus, P_r_ emerges as a key presynaptic target through which distinct molecular sleep regulators mediate their effects on sleep behavior.

## Discussion

The Synaptic Homeostasis Hypothesis (SHY) suggests the net synaptic strength is the central parameter that is modulated by sleep and wakefulness (*1*). Yet directly measuring net synaptic strength across all excitatory synapses in the brain by electrophysiology remains technically challenging, making SHY difficult to evaluate. However, our functional assessment from two anatomically defined excitatory synapses in *Drosophila* indicates that the synaptic strength of these two circuits remained stable or even decreased following wakefulness, raising questions about SHY’s validity, at least in flies. In contrast, presynaptic release probability (P_r_) consistently declined after extended wakefulness and was restored by sleep at multiple synapses in both flies and mice. Additionally, multiple known sleep-regulatory signaling pathways converge on P_r_, rather than on synaptic strength. Thus, based on these functional assessments, we propose an alternative model: the Presynaptic Release Probability Homeostasis Hypothesis (P_r_HY, pronounced “phi”). During wakefulness, ongoing information processing demands sustained synaptic transmission, placing a continuous load on the presynaptic release machinery. Over time, this sustained activity leads to a gradual decline in P_r_, representing a form of “fatigue” at presynaptic terminals that impairs the efficiency and temporal precision of neurotransmission. Sleep functions to restore P_r_, likely by enhancing the efficiency of synaptic vesicle release, thereby reestablishing effective and reliable synaptic transmission.

P_r_HY is compatible with existing observations regarding sleep-dependent synaptic changes. Several studies in rodents report an increased frequency of miniature excitatory postsynaptic currents (mEPSCs) in the cortex after sleep loss (*18*, *60*, *61*), with few exceptions (*19*, *62*). In our own recordings from pyramidal neurons in the mouse mPFC and BC, we observed similar sleep-loss-induced increases in mEPSC frequency, without accompanying changes in amplitude (fig. S8). Given our concurrent observations of reduced P_r_ after sleep deprivation, this increased mEPSC frequency likely reflects an elevated number of functional synapses. This interpretation aligns with prior studies that report increased AMPA receptor expression in the cortex following prolonged wakefulness (*10*, *12*), further indicating an increase in the number of functional synapses. Together, these observations may represent a form of homeostatic synaptic plasticity that occurs during wakefulness in rodents, in which the number of functional synapses increases to counterbalance the reduced P_r_. Notably, this form of homeostatic compensation during wakefulness appears not to exist in every circuit in the *Drosophila* brain, as GRASP (GFP□Reconstitution□Across□Synaptic□Partners) analyses suggest circuit-specific changes in synapse number after sleep deprivation (*13*). Instead, compensation for reduced P_r_ may occur presynaptically in flies, as suggested by a widespread increase of active zone scaffold protein Bruchpilot (Brp) across excitatory synapses following sleep loss (*11*). Indeed, knockdown of the voltage-gated Ca^2+^ channel Cac, which reduces P_r_, also leads to increased Brp expression in fly synapses (*63*). This compensatory Brp upregulation during wakefulness may also actively contribute to sleep-wake regulation (*64*), either by further reducing P_r_ or by engaging in other unidentified presynaptic processes. Collectively, these molecular, structural, and mEPSC synaptic changes likely reflect an evolutionarily conserved compensatory mechanism to counteract wakefulness-induced reductions in P_r_. It also suggests that total synaptic transmission or synaptic strength in many circuits is promptly maintained during wakefulness or irrespective of sleep-wake states, while P_r_ is the key synaptic parameter regulated by sleep and wakefulness.

The observation that sleep deprivation reduces P_r_ at excitatory synapses in both *Drosophila* and mammalian cortex highlights P_r_ as an evolutionarily conserved mechanism in sleep regulation. Similar reductions in Pr have also been reported in the medial prefrontal cortex– to–nucleus accumbens pathway (*65*), suggesting that wakefulness-induced decreases in P_r_ are a general feature of cortical excitatory synapses. Genetic evidence also aligns with this model: a hypomorphic mutation in the mouse *VAMP2* gene, the homolog of *Drosophila nSyb*, reduces P_r_ (*66*) and leads to persistently elevated slow-wave activity (SWA) (*67*), a well-accepted marker of sleep pressure in mammals. This parallels our finding that nSyb knockdown in flies increases sleep need, reinforcing the idea that P_r_ serves as an evolutionarily conserved mechanism for encoding and regulating sleep homeostasis.

We demonstrate that multiple known sleep-regulatory signaling pathways converge specifically on P_r_ rather than synaptic strength. These findings connect sleep-regulatory kinases and neuronal redox states to the notion that sleep need is tightly linked to synaptic function. Moreover, they provide a potential mechanistic explanation for how wakefulness reduces P_r_. During wakefulness, neuronal activity, such as that associated with learning, may influence cyclic AMP (cAMP) signaling pathways (*68*), thereby altering the activities of sleep-regulating kinases, including PKA and Sik3. Changes in the activities of these kinases subsequently reduce P_r_. Concurrently, wakefulness-induced neuronal activity elevates cellular stress, particularly oxidative stress (*57*, *69*), also contributing to reduced P_r_. Thus, P_r_ may integrate various neuronal signals induced by wakefulness, serving as a presynaptic mechanism that determines sleep need.

Our data indicates that the reduction in P_r_ during wakefulness stems from impaired Ca^2+^-dependent vesicle release. Specifically, we observed either diminished Ca^2+^ influx or weakened coupling between voltage-gated Ca^2+^ channels and synaptic vesicles. Although the precise mechanisms may differ across circuits, these changes converge on a shared outcome: reduced efficiency of vesicle exocytosis. These findings highlight the value of direct, functional measurements in revealing how sleep–wake states influence synaptic transmission. Notably, a recent study using human cortical slices showed that mimicking slow-wave activity enhances P_r_ by broadening axonal action potentials and enhancing Ca^2+^ influx at presynaptic terminals (*70*). This supports P_r_HY that sleep reverses wake-induced reduction of P_r_ by restoring the machinery responsible for Ca^2+^-dependent synaptic vesicle release. Thus, despite mechanistic differences across synapses, sleep appears to promote a conserved recovery of P_r_ through Ca^2+^-dependent processes.

In summary, P_r_HY provides a functional, mechanistic, and evolutionarily conserved framework for understanding how sleep modulates synaptic function. By identifying presynaptic release probability (P_r_) as a key parameter regulated by sleep and wakefulness, P_r_HY shifts the focus from synaptic strength to the efficiency of neurotransmission. Our cross-species analyses, combining electrophysiology, imaging, and ultrastructural data, support this model and highlight the value of direct functional assessment. Future studies should examine whether additional synaptic processes, including vesicle recycling, endocytosis, or postsynaptic mechanisms, contribute to sleep need alongside changes in P_r_. It will also be important to determine how wake-induced P_r_ reduction impacts sleep-dependent cognitive processes and behavior, particularly through alterations in short-term synaptic plasticity.

## Acknowledgements

We thank Drs. Jason Rihel, Jakob B. Sørensen, Jianyuan Sun, and Mark Wu, for their helpful comments on the manuscript. We thank the Bloomington *Drosophila* Stock Center, Drs. Kate O’Connor-Giles, Nicolas Tapon, and Mark Wu, for providing the fly stocks. We also thank the Service Laboratory of Electron Microscopy at the Institute of Molecular Genetics of the Czech Academy of Sciences for performing tomography. This research was supported by the European Research Council (StG 758580 to S.L.), Fonds voor Wetenschappelijk Onderzoek (FWO) Research Project (G052922N to S.L.), National Institutes of Health (NS091546 and NS126654 to D.D.)

## Author Contributions

Y.W. and S. Liu conceived the project. Y.W. performed most adult fly electrophysiology experiments with help from Y.-C.H. K.W. performed mouse electrophysiology experiments. Y.W., F.L., and M.H. performed mouse behavioral experiments. Y.W., Y.-C.H., and C.-C.C. performed functional imaging experiments. Y.W., K.V., F.L., and N.V.G. performed electron microscopy experiments. Y.W. and S. Loomba analyzed electron microscopy data. Y.W. and F.L. prepared fly genetics with help from M.C.D. Y.W. performed fly behavioral experiments with help from E.-S.B. V.U. performed fly NMJ experiments with input from P.V. S.H., C.P., and S.S. provided *unc13A* fly lines with initial characterization of their sleep behavior. J.C. and D.D. developed and characterized the *20xUAS-Syt-mScarlet-GCaMP8m* flies. S.B. and L.L. synthesized PA-Nic2 reagent. Y.W. and S. Liu wrote the manuscript with feedback from all authors.

## Competing interests

The authors have no competing interests to declare.

## Data and materials availability

The data supporting the findings of this study are available within the article and its Supplementary Materials. Raw datasets generated during the current study are available from the corresponding author upon request. Source data will be provided with publication.

## Supplementary Materials

Materials and Methods

Figs. S1 to S9

Tables S1 to S4

References

## Notes

### Competing Interest Statement

The authors have declared no competing interest.

## References and Notes

1. G. Tononi, C. Cirelli, Sleep function and synaptic homeostasis. Sleep Med. Rev. 10, 49– 62 (2006).

2. M. G. Frank, The mystery of sleep function: current perspectives and future directions. Rev. Neurosci. 17, 375–392 (2006).

3. J. Seibt, M. G. Frank, Primed to sleep: The dynamics of synaptic plasticity across brain states. Front. Syst. Neurosci. 13, 2 (2019).

4. C. Cirelli, G. Tononi, Linking the need to sleep with synaptic function, Science (New York, N.Y.). 366 (2019)pp. 189–190.

5. L. de Vivo, M. Bellesi, W. Marshall, E. A. Bushong, M. H. Ellisman, G. Tononi, C. Cirelli, Ultrastructural evidence for synaptic scaling across the wake/sleep cycle. Science 355, 507–510 (2017).

6. R. Havekes, A. J. Park, J. C. Tudor, V. G. Luczak, R. T. Hansen, S. L. Ferri, V. M. Bruinenberg, S. G. Poplawski, J. P. Day, S. J. Aton, K. Radwańska, P. Meerlo, M. D. Houslay, G. S. Baillie, T. Abel, Sleep deprivation causes memory deficits by negatively impacting neuronal connectivity in hippocampal area CA1. Elife 5 (2016).

7. D. Bushey, G. Tononi, C. Cirelli, Sleep and synaptic homeostasis: structural evidence in Drosophila. Science 332, 1576–1581 (2011).

8. G. Yang, C. S. W. Lai, J. Cichon, L. Ma, W. Li, W.-B. Gan, Sleep promotes branch-specific formation of dendritic spines after learning. Science 344, 1173–1178 (2014).

9. C. Cirelli, C. M. Gutierrez, G. Tononi, Extensive and divergent effects of sleep and wakefulness on brain gene expression. Neuron 41, 35–43 (2004).

10. V. V. Vyazovskiy, C. Cirelli, M. Pfister-Genskow, U. Faraguna, G. Tononi, Molecular and electrophysiological evidence for net synaptic potentiation in wake and depression in sleep. Nat. Neurosci. 11, 200–208 (2008).

11. G. F. Gilestro, G. Tononi, C. Cirelli, Widespread changes in synaptic markers as a function of sleep and wakefulness in Drosophila. Science 324, 109–112 (2009).

12. G. H. Diering, R. S. Nirujogi, R. H. Roth, P. F. Worley, A. Pandey, R. L. Huganir, Homer1a drives homeostatic scaling-down of excitatory synapses during sleep. Science 355, 511–515 (2017).

13. J. T. Weiss, J. M. Donlea, Sleep deprivation results in diverse patterns of synaptic scaling across the Drosophila mushroom bodies. Curr. Biol. 31, 3248–3261.e3 (2021).

14. M. Matsuzaki, G. C. Ellis-Davies, T. Nemoto, Y. Miyashita, M. Iino, H. Kasai, Dendritic spine geometry is critical for AMPA receptor expression in hippocampal CA1 pyramidal neurons. Nat. Neurosci. 4, 1086–1092 (2001).

15. M. Matsuzaki, N. Honkura, G. C. R. Ellis-Davies, H. Kasai, Structural basis of long-term potentiation in single dendritic spines. Nature 429, 761–766 (2004).

16. A. Thomazeau, M. Bosch, S. Essayan-Perez, S. A. Barnes, H. De Jesus-Cortes, M. F. Bear, Dissociation of functional and structural plasticity of dendritic spines during NMDAR and mGluR-dependent long-term synaptic depression in wild-type and fragile X model mice. Mol. Psychiatry 26, 4652–4669 (2021).

17. D. Choquet, Shifting rules in a brain disorder, Science (New York, N.Y.). 383 (2024)pp. 950–951.

18. Z.-W. Liu, U. Faraguna, C. Cirelli, G. Tononi, X.-B. Gao, Direct evidence for wake-related increases and sleep-related decreases in synaptic strength in rodent cortex. J. Neurosci. 30, 8671–8675 (2010).

19. B. A. Cary, G. G. Turrigiano, Stability of neocortical synapses across sleep and wake states during the critical period in rats. Elife 10, e66304 (2021).

20. S. Dorkenwald, A. Matsliah, A. R. Sterling, P. Schlegel, S.-C. Yu, C. E. McKellar, A. Lin, M. Costa, K. Eichler, Y. Yin, W. Silversmith, C. Schneider-Mizell, C. S. Jordan, D. Brittain, A. Halageri, K. Kuehner, O. Ogedengbe, R. Morey, J. Gager, K. Kruk, E. Perlman, R. Yang, D. Deutsch, D. Bland, M. Sorek, R. Lu, T. Macrina, K. Lee, J. A. Bae, S. Mu, B. Nehoran, E. Mitchell, S. Popovych, J. Wu, Z. Jia, M. A. Castro, N. Kemnitz, D. Ih, A. S. Bates, N. Eckstein, J. Funke, F. Collman, D. D. Bock, G. S. X. E. Jefferis, H. S. Seung, M. Murthy, FlyWire Consortium, Neuronal wiring diagram of an adult brain. Nature 634, 124–138 (2024).

21. P. Schlegel, Y. Yin, A. S. Bates, S. Dorkenwald, K. Eichler, P. Brooks, D. S. Han, M. Gkantia, M. Dos Santos, E. J. Munnelly, G. Badalamente, L. Serratosa Capdevila, V. A. Sane, A. M. C. Fragniere, L. Kiassat, M. W. Pleijzier, T. Stürner, I. F. M. Tamimi, C. R. Dunne, I. Salgarella, A. Javier, S. Fang, E. Perlman, T. Kazimiers, S. R. Jagannathan, A. Matsliah, A. R. Sterling, S.-C. Yu, C. E. McKellar, FlyWire Consortium, M. Costa, H. S. Seung, M. Murthy, V. Hartenstein, D. D. Bock, G. S. X. E. Jefferis, Whole-brain annotation and multi-connectome cell typing of Drosophila. Nature 634, 139–152 (2024).

22. R. I. Wilson, G. Laurent, Role of GABAergic inhibition in shaping odor-evoked spatiotemporal patterns in the Drosophila antennal lobe. J. Neurosci. 25, 9069–9079 (2005).

23. K. I. Nagel, E. J. Hong, R. I. Wilson, Synaptic and circuit mechanisms promoting broadband transmission of olfactory stimulus dynamics. Nat. Neurosci. 18, 56–65 (2015).

24. T. Hige, Y. Aso, M. N. Modi, G. M. Rubin, G. C. Turner, Heterosynaptic plasticity underlies aversive olfactory learning in Drosophila. Neuron 88, 985–998 (2015).

25. N. C. Klapoetke, Y. Murata, S. S. Kim, S. R. Pulver, A. Birdsey-Benson, Y. K. Cho, T. K. Morimoto, A. S. Chuong, E. J. Carpenter, Z. Tian, J. Wang, Y. Xie, Z. Yan, Y. Zhang, B. Y. Chow, B. Surek, M. Melkonian, V. Jayaraman, M. Constantine-Paton, G. K.-S. Wong, E. S. Boyden, Independent optical excitation of distinct neural populations. Nat. Methods 11, 338–346 (2014).

26. L. E. Dobrunz, C. F. Stevens, Heterogeneity of release probability, facilitation, and depletion at central synapses. Neuron 18, 995–1008 (1997).

27. T. Sawada, Y. Iino, K. Yoshida, H. Okazaki, S. Nomura, C. Shimizu, T. Arima, M. Juichi, S. Zhou, N. Kurabayashi, T. Sakurai, S. Yagishita, M. Yanagisawa, T. Toyoizumi, H. Kasai, S. Shi, Prefrontal synaptic regulation of homeostatic sleep pressure revealed through synaptic chemogenetics. Science 385, 1459–1465 (2024).

28. S. Banala, X.-T. Jin, T. L. Dilan, S.-H. Sheu, D. E. Clapham, R. M. Drenan, L. D. Lavis, Elucidating and optimizing the photochemical mechanism of coumarin-caged tertiary amines. J. Am. Chem. Soc. 146, 20627–20635 (2024).

29. I. Bucurenciu, A. Kulik, B. Schwaller, M. Frotscher, P. Jonas, Nanodomain coupling between Ca2+ channels and Ca2+ sensors promotes fast and efficient transmitter release at a cortical GABAergic synapse. Neuron 57, 536–545 (2008).

30. N. Rebola, M. Reva, T. Kirizs, M. Szoboszlay, A. Lőrincz, G. Moneron, Z. Nusser, D. A. DiGregorio, Distinct nanoscale calcium channel and synaptic vesicle topographies contribute to the diversity of synaptic function. Neuron 109, 3178 (2021).

31. M. A. Böhme, C. Beis, S. Reddy-Alla, E. Reynolds, M. M. Mampell, A. T. Grasskamp, J. Lützkendorf, D. D. Bergeron, J. H. Driller, H. Babikir, F. Göttfert, I. M. Robinson, C. J. O’Kane, S. W. Hell, M. C. Wahl, U. Stelzl, B. Loll, A. M. Walter, S. J. Sigrist, Active zone scaffolds differentially accumulate Unc13 isoforms to tune Ca(2+) channel-vesicle coupling. Nat. Neurosci. 19, 1311–1320 (2016).

32. R. J. Kittel, C. Wichmann, T. M. Rasse, W. Fouquet, M. Schmidt, A. Schmid, D. A. Wagh, C. Pawlu, R. R. Kellner, K. I. Willig, S. W. Hell, E. Buchner, M. Heckmann, S. J. Sigrist, Bruchpilot promotes active zone assembly, Ca2+ channel clustering, and vesicle release. Science 312, 1051–1054 (2006).

33. K. M. Lembke, A. D. Law, J. Ahrar, D. B. Morton, Deletion of a specific exon in the voltage-gated calcium channel gene cacophony disrupts locomotion in Drosophila larvae. J. Exp. Biol. 222 (2019).

34. J. J. Bruckner, S. J. Gratz, J. K. Slind, R. R. Geske, A. M. Cummings, S. E. Galindo, L. K. Donohue, K. M. O’Connor-Giles, Fife, a Drosophila Piccolo-RIM homolog, promotes active zone organization and neurotransmitter release. J. Neurosci. 32, 17048–17058 (2012).

35. P. S. Kaeser, L. Deng, Y. Wang, I. Dulubova, X. Liu, J. Rizo, T. C. Südhof, RIM proteins tether Ca2+ channels to presynaptic active zones via a direct PDZ-domain interaction. Cell 144, 282–295 (2011).

36. E. R. Graf, V. Valakh, C. M. Wright, C. Wu, Z. Liu, Y. Q. Zhang, A. DiAntonio, RIM promotes calcium channel accumulation at active zones of the Drosophila neuromuscular junction. J. Neurosci. 32, 16586–16596 (2012).

37. J. J. Bruckner, H. Zhan, S. J. Gratz, M. Rao, F. Ukken, G. Zilberg, K. M. O’Connor-Giles, Fife organizes synaptic vesicles and calcium channels for high-probability neurotransmitter release. J. Cell Biol. 216, 231–246 (2017).

38. B. Deng, Q. Li, X. Liu, Y. Cao, B. Li, Y. Qian, R. Xu, R. Mao, E. Zhou, W. Zhang, J. Huang, Y. Rao, Chemoconnectomics: Mapping chemical transmission in Drosophila. Neuron 101, 876–893.e4 (2019).

39. T. Osterwalder, K. S. Yoon, B. H. White, H. Keshishian, A conditional tissue-specific transgene expression system using inducible GAL4. Proc. Natl. Acad. Sci. U. S. A. 98, 12596–12601 (2001).

40. N. Brose, A. G. Petrenko, T. C. Südhof, R. Jahn, Synaptotagmin: a calcium sensor on the synaptic vesicle surface. Science 256, 1021–1025 (1992).

41. J. T. Littleton, M. Stern, K. Schulze, M. Perin, H. J. Bellen, Mutational analysis of Drosophila synaptotagmin demonstrates its essential role in Ca(2+)-activated neurotransmitter release. Cell 74, 1125–1134 (1993).

42. A. DiAntonio, R. W. Burgess, A. C. Chin, D. L. Deitcher, R. H. Scheller, T. L. Schwarz, Identification and characterization of Drosophila genes for synaptic vesicle proteins. J. Neurosci. 13, 4924–4935 (1993).

43. H. T. McMahon, M. Missler, C. Li, T. C. Südhof, Complexins: cytosolic proteins that regulate SNAP receptor function. Cell 83, 111–119 (1995).

44. S. Huntwork, J. T. Littleton, A complexin fusion clamp regulates spontaneous neurotransmitter release and synaptic growth. Nat. Neurosci. 10, 1235–1237 (2007).

45. M. Müller, E. C. G. Pym, A. Tong, G. W. Davis, Rab3-GAP controls the progression of synaptic homeostasis at a late stage of vesicle release. Neuron 69, 749–762 (2011).

46. J. E. Richmond, W. S. Davis, E. M. Jorgensen, UNC-13 is required for synaptic vesicle fusion in C. elegans. Nat. Neurosci. 2, 959–964 (1999).

47. I. Augustin, C. Rosenmund, T. C. Südhof, N. Brose, Munc13-1 is essential for fusion competence of glutamatergic synaptic vesicles. Nature 400, 457–461 (1999).

48. B. Aravamudan, T. Fergestad, W. S. Davis, C. K. Rodesch, K. Broadie, Drosophila UNC-13 is essential for synaptic transmission. Nat. Neurosci. 2, 965–971 (1999).

49. V. Budnik, Y. H. Koh, B. Guan, B. Hartmann, C. Hough, D. Woods, M. Gorczyca, Regulation of synapse structure and function by the Drosophila tumor suppressor gene dlg. Neuron 17, 627–640 (1996).

50. M. Jusyte, N. Blaum, M. A. Böhme, M. M. M. Berns, A. E. Bonard, Á. B. Vámosi, K. V. Pushpalatha, J. R. L. Kobbersmed, A. M. Walter, Unc13A dynamically stabilizes vesicle priming at synaptic release sites for short-term facilitation and homeostatic potentiation. Cell Rep. 42, 112541 (2023).

51. H. Funato, C. Miyoshi, T. Fujiyama, T. Kanda, M. Sato, Z. Wang, J. Ma, S. Nakane, J. Tomita, A. Ikkyu, M. Kakizaki, N. Hotta-Hirashima, S. Kanno, H. Komiya, F. Asano, T. Honda, S. J. Kim, K. Harano, H. Muramoto, T. Yonezawa, S. Mizuno, S. Miyazaki, L. Connor, V. Kumar, I. Miura, T. Suzuki, A. Watanabe, M. Abe, F. Sugiyama, S. Takahashi, K. Sakimura, Y. Hayashi, Q. Liu, K. Kume, S. Wakana, J. S. Takahashi, M. Yanagisawa, Forward-genetics analysis of sleep in randomly mutagenized mice. Nature 539, 378–383 (2016).

52. Z. Wang, J. Ma, C. Miyoshi, Y. Li, M. Sato, Y. Ogawa, T. Lou, C. Ma, X. Gao, C. Lee, T. Fujiyama, X. Yang, S. Zhou, N. Hotta-Hirashima, D. Klewe-Nebenius, A. Ikkyu, M. Kakizaki, S. Kanno, L. Cao, S. Takahashi, J. Peng, Y. Yu, H. Funato, M. Yanagisawa, Q. Liu, Quantitative phosphoproteomic analysis of the molecular substrates of sleep need. Nature 558, 435–439 (2018).

53. M. N. Wu, K. Ho, A. Crocker, Z. Yue, K. Koh, A. Sehgal, The effects of caffeine on sleep in Drosophila require PKA activity, but not the adenosine receptor. J. Neurosci. 29, 11029–11037 (2009).

54. V. M. Hill, R. M. O’Connor, G. B. Sissoko, I. S. Irobunda, S. Leong, J. C. Canman, N. Stavropoulos, M. Shirasu-Hiza, A bidirectional relationship between sleep and oxidative stress in Drosophila. PLoS Biol. 16, e2005206 (2018).

55. P. R. Haynes, E. S. Pyfrom, Y. Li, C. Stein, V. A. Cuddapah, J. A. Jacobs, Z. Yue, A. Sehgal, A neuron-glia lipid metabolic cycle couples daily sleep to mitochondrial homeostasis. Nat. Neurosci. 27, 666–678 (2024).

56. H. O. Rorsman, M. A. Müller, P. Z. Liu, L. G. Sanchez, A. Kempf, S. Gerbig, B. Spengler, G. Miesenböck, Sleep pressure accumulates in a voltage-gated lipid peroxidation memory. Nature 641, 232–239 (2025).

57. Y. Tian, L. Kang, N. T. Ha, J. Deng, D. Liu, Hydrogen peroxide in midbrain sleep neurons regulates sleep homeostasis. Cell Metab. 37, 1442–1452.e7 (2025).

58. E. Reimund, The free radical flux theory of sleep. Med. Hypotheses 43, 231–233 (1994).

59. T. L. Parkes, A. J. Elia, D. Dickinson, A. J. Hilliker, J. P. Phillips, G. L. Boulianne, Extension of Drosophila lifespan by overexpression of human SOD1 in motorneurons. Nat. Genet. 19, 171–174 (1998).

60. M. C. D. Bridi, F.-J. Zong, X. Min, N. Luo, T. Tran, J. Qiu, D. Severin, X.-T. Zhang, G. Wang, Z.-J. Zhu, K.-W. He, A. Kirkwood, Daily Oscillation of the Excitation-Inhibition Balance in Visual Cortical Circuits. Neuron 105, 621–629.e4 (2020).

61. S. Hatori, S. T. Yamaguchi, R. Kobayashi, K. Okamoto, Z. Zhou, K. T. Kotake, F. Matsui, H. Hioki, H. Norimoto, Sleep homeostasis in lizards and the role of the cortex. Proc. Natl. Acad. Sci. U. S. A. 122, e2415929122 (2025).

62. B. D. Winters, Y. H. Huang, Y. Dong, J. M. Krueger, Sleep loss alters synaptic and intrinsic neuronal properties in mouse prefrontal cortex. Brain Res. 1420, 1–7 (2011).

63. E. Rozenfeld, N. Ehmann, J. E. Manoim, R. J. Kittel, M. Parnas, Homeostatic synaptic plasticity rescues neural coding reliability. Nat. Commun. 14, 2993 (2023).

64. S. Huang, C. Piao, C. B. Beuschel, T. Götz, S. J. Sigrist, Presynaptic active zone plasticity encodes sleep need in Drosophila. Curr. Biol. 30, 1077–1091.e5 (2020).

65. Z. Liu, Y. Wang, L. Cai, Y. Li, B. Chen, Y. Dong, Y. H. Huang, Prefrontal cortex to accumbens projections in sleep regulation of reward. J. Neurosci. 36, 7897–7910 (2016).

66. G. T. Banks, M. C. C. Guillaumin, I. Heise, P. Lau, M. Yin, N. Bourbia, C. Aguilar, M. R. Bowl, C. Esapa, L. A. Brown, S. Hasan, E. Tagliatti, E. Nicholson, R. S. Bains, S. Wells, V. V. Vyazovskiy, K. Volynski, S. N. Peirson, P. M. Nolan, Forward genetics identifies a novel sleep mutant with sleep state inertia and REM sleep deficits. Sci. Adv. 6, eabb3567 (2020).

67. M. C. C. Guillaumin, C. D. Harding, L. B. Krone, T. Yamagata, M. C. Kahn, C. Blanco-Duque, G. T. Banks, P. Achermann, C. Diniz Behn, P. M. Nolan, S. N. Peirson, V. V. Vyazovskiy, Deficient synaptic neurotransmission results in a persistent sleep-like cortical activity across vigilance states in mice. Curr. Biol. 35, 1716–1729.e3 (2025).

68. M. Brunelli, V. Castellucci, E. R. Kandel, Synaptic facilitation and behavioral sensitization in Aplysia: possible role of serotonin and cyclic AMP. Science 194, 1178– 1181 (1976).

69. Y. Rabah, J.-P. Berwick, N. Sagar, L. Pasquer, P.-Y. Plaçais, T. Preat, Astrocyte-to-neuron H2O2 signalling supports long-term memory formation in Drosophila and is impaired in an Alzheimer’s disease model. Nat. Metab. 7, 321–335 (2025).

70. F. X. Mittermaier, T. Kalbhenn, R. Xu, J. Onken, K. Faust, T. Sauvigny, U. W. Thomale, A. M. Kaindl, M. Holtkamp, S. Grosser, P. Fidzinski, M. Simon, H. Alle, J. R. P. Geiger, Membrane potential states gate synaptic consolidation in human neocortical tissue. Nat. Commun. 15, 10340 (2024).

71. Q. Geissmann, L. Garcia Rodriguez, E. J. Beckwith, A. S. French, A. R. Jamasb, G. F. Gilestro, Ethoscopes: An open platform for high-throughput ethomics. PLoS Biol. 15, e2003026 (2017).

72. Q. Geissmann, L. Garcia Rodriguez, E. J. Beckwith, G. F. Gilestro, Rethomics: An R framework to analyse high-throughput behavioural data. PLoS One 14, e0209331 (2019).

73. J. Dopp, A. Ortega, K. Davie, S. Poovathingal, E.-S. Baz, S. Liu, Single-cell transcriptomics reveals that glial cells integrate homeostatic and circadian processes to drive sleep-wake cycles. Nat. Neurosci. 27, 359–372 (2024).

74. M. M. Hoekstra, M. Jan, G. Katsioudi, Y. Emmenegger, P. Franken, The sleep-wake distribution contributes to the peripheral rhythms in PERIOD-2. Elife 10 (2021).

75. P. Thévenaz, U. E. Ruttimann, M. Unser, A pyramid approach to subpixel registration based on intensity. IEEE Trans. Image Process. 7, 27–41 (1998).

76. K. Vints, P. Baatsen, N. V. Gounko, An accelerated procedure for approaching and imaging of optically branded region of interest in tissue. Methods Cell Biol. 162, 205– 221 (2021).

77. D. N. Mastronarde, Automated electron microscope tomography using robust prediction of specimen movements. J. Struct. Biol. 152, 36–51 (2005).

78. J. R. Kremer, D. N. Mastronarde, J. R. McIntosh, Computer visualization of three-dimensional image data using IMOD. J. Struct. Biol. 116, 71–76 (1996).

79. K. M. Boergens, M. Berning, T. Bocklisch, D. Bräunlein, F. Drawitsch, J. Frohnhofen, T. Herold, P. Otto, N. Rzepka, T. Werkmeister, D. Werner, G. Wiese, H. Wissler, M. Helmstaedter, webKnossos: efficient online 3D data annotation for connectomics. Nat. Methods 14, 691–694 (2017).

80. P. S. Kaeser, W. G. Regehr, The readily releasable pool of synaptic vesicles. Curr. Opin. Neurobiol. 43, 63–70 (2017).

