## Supplementary Materials for "Presynaptic Release Probability Determines the Need for Sleep"

Yifan Wu *et al.*

**The file includes:**

Materials and Methods

Figs. S1 to S9

Tables S1 to S4

References

Materials and Methods

***Drosophila* strains and husbandry**

*Drosophila melanogaster* were reared at 25 °C and 60% relative humidity on standard cornmeal fly food under a 12:12-hour light-dark cycle. All experiments were performed on randomly selected mated females aged 4–10 days post eclosion. The following fly lines were obtained from the Vienna *Drosophila* Resource Center: *VT064246-Gal4* (204311), *UAS-Syt1-RNAi* (100608), *UAS-nSyb-RNAi* (104531), *UAS-cpx-RNAi* (21477), *UAS-Rab3-GAP-RNAi* (106905), *UAS-Fife-RNAi* (11059), *UAS-Rim-RNAi* (39384, used in Fig. 3c,d), and *UAS-dlg1-RNAi* (41134). Additional lines from the Bloomington *Drosophila* Stock Center (BDSC) included *Mz19-Gal4* (34497), *13XLexAop2-Chronos-mVenus-p10* (77113), *UAS-CD4-tdTom* (35841), *20XUAS-CsChrimson-mCherry* (82181), *Or88a-Gal4* (23294), *Mz19-QF* (41573), *10XQUAS-6XGFP* (52264), *VA1d-adPN-split-Gal4* (87516), *ChAT-T2A-Gal4* (84618), *R13F02-lexA(JK22c)* (94676), *ELAV-GS* (43642), *cac^Exon7∆^* (93122), *Rim^Ex73^* (78046), *UAS-Dcr-2* (24650 and 24651), *UAS-Rim-RNAi* (4454, used in Fig. 3f,g), and *UAS-hSOD1* (33606). Additional lines were generated in this study or obtained from collaborators: *10XUAS-GFP-p10 (JK22c)* (generated in this study); *20XUAS-Syt-mScarlet-GCaMP8m* (from D.K. Dickman, University of Southern California); *Fife^AC^* (from K.M. O’Connor-Giles, Brown University,); *Unc13A^WT^*, *Unc13A^CaMWRWR^*, and *UAS-Unc13-RNAi* (from S.J. Sigrist, Berlin Free University); *UAS-Sik3^S563A^* (from N. Tapon, Francis Crick Institute), and *UAS-PKAR* (from M.N. Wu, Johns Hopkins University). All lines were outcrossed into the wild-type *iso^31^* (BDSC5905) background at least 5 times, except the few lines for GeneSwitch induced RNAi knockdown experiments. The exact genotypes and genetic background of fly lines used in each figure are summarized in Table S1 and S3.

For GeneSwitch-related experiments, flies were fed either 500 μM RU486 (dissolved in 0.4% ethanol) or 0.4% ethanol alone, starting 4–6 days after eclosion. Both solutions were prepared in 2% agarose/5% sucrose tubes. In all RNAi-related experiments, *UAS*-*Dcr*-2 was used to enhance RNAi efficiency.

For optogenetic experiments, one-day-old flies were placed on standard food supplemented with 400 μM all-trans-retinal and kept for four days to ensure adequate expression and activation of Chronos or Chrimson in targeted neurons.

**Mice**

Male C57BL/6 mice, aged 9 weeks, were kept under controlled ambient conditions (20–23 °C, 40–60% humidity) with a 12:12-hour light-dark cycle and provided with unrestricted access to food and water. Prior to the experiments, mice were individually housed and re-entrained to the target 12:12-hour light-dark cycle (light-on at 6am) for a minimum of 10 days. All experiments were conducted in compliance with KU Leuven ethical standards and received approval from the KU Leuven Ethical Committee for Animal Experimentation.

**Sleep behaviour in flies**

Mated female flies (4–10 days old) were entrained to a 12:12-hour light-dark cycle for at least 3 days before being transferred into 65 mm × 5 mm glass locomotor tubes containing 5% sucrose and 2% agar food (supplemented with RU-486 for GeneSwitch experiments) for sleep monitoring. Sleep data were recorded using either *Drosophila* Activity Monitoring (DAM) Systems (Trikinetics) with 20-second bins or modified Ethoscopes system (*71*) with 1 second bins. Sleep parameters, including sleep amount, bout number, and bout length, were analysed over 12- or 24-hour periods and averaged across two or more days using a custom R script based on the Rethomics (*72*) package. Waking activity was calculated as the mean distance travelled between consecutive frames during wake periods, based on images acquired every 10 seconds.

For sleep deprivation, flies in the DAM system were subjected to randomized 1.5-second shaking per minute using a Trikinetics vortexer. Flies monitored with Ethoscopes were deprived of sleep using a custom designed rotational module (*71*, *73*), trigged after 10 seconds of fly immobility. Recovery sleep was measured during the 4-hour period following a 12-hour sleep deprivation from zeitgeber time 12 (ZT12) to ZT24. Recovery sleep was calculated as the difference between the sleep obtained during this window (ZT0–4) and the corresponding circadian period on the preceding day. The sleep monitoring and deprivation methods used for each experiment are summarized in Table S1 and S3.

For the mechanical arousal experiment, flies received either a 0.3-second weak stimulus at the lowest vortex intensity or a 3-second strong stimulus at intensity level 3, using the Trikinetics vortexer at ZT16. Flies that remained inactive for at least 5 minutes before stimulation were considered asleep, while those that moved within 3 minutes after the stimulus were classified as “aroused.”

**Sleep deprivation in mice**

In mice, sleep deprivation was conducted between ZT0 and ZT6 using a gentle handling protocol, as previously described (*74*). Mice were continuously monitored and briefly stimulated upon the appearance of behavioural signs of sleep, using mild, non-invasive interventions such as removing paper tissue or cage litter, gently tapping the cage, or moving a pipette near the animal. Control mice remained undisturbed in their home cages throughout the same period.

**Electrophysiological recordings**

For whole-cell patch-clamp recordings in adult female flies, aged 5–8 days post-eclosion were used. Prior to dissection, flies were anesthetized on ice for ~30 seconds. Brains were then dissected in the external saline containing (in mM): 103 NaCl, 3 KCl, 5 TES, 8 trehalose, 10 glucose, 26 NaHCO3, 1 NaH_2_PO_4_, 1.5 CaCl_2_, and 4 MgCl_2_ (osmolarity adjusted to 270–275 mOsm), bubbled with 95% O_2_ and 5% CO_2_ to a final pH of 7.3. Brains were then treated with an enzymatic cocktail, collagenase (0.2 mg/mL), protease XIV (0.4 mg/mL), and dispase (0.6 mg/mL) (Sigma-Aldrich) for 90 seconds at 22 °C to expose and clean the surface of targeted neurons. Recordings were performed either at zeitgeber time 0 (ZT0) and continued for up to 3 hours, or at the specific times indicated in the corresponding figure panels. Targeted neurons were visualized by either GFP or tdTomato fluorescence and infrared-differential interference contrast (IR-DIC) optics using a fixed-stage upright microscope (Scientifica SliceScope). During recordings, the saline was constantly perfused to the recording chamber at 2 ml/min. The patch-clamp pipette (4-5 MΩ) was filled with the internal solution containing (in mM): 140 Cs-Gluconate, 10 HEPES, 1 EGTA, 4 MgATP, 0.5 Na₃GTP, 1 KCl, 13 biocytin hydrazide, and 10 QX-314 Chloride (pH 7.2, ~270 mOsm). For current-clamp recordings, Cs-gluconate was replaced with K-gluconate. Signals were gathered with a Multiclamp 700B amplifier (Molecular Devices), filtered at 3 kHz and then digitized at 20 kHz via Digidata 1550B interface (Molecular Devices). In current-clamp mode, bridge balance was used to compensate for series resistance. To assess intrinsic excitability in VA1d PNs, a series of 1-s current injections (−80 to +140 pA in 20-pA steps) was delivered from a membrane potential of −60 mV. Input resistance was calculated from the mean steady-state voltage response to a series of 250-ms hyperpolarising current injections (−10 pA) delivered prior to the step protocol. Firing frequency was quantified as the number of action potentials elicited during each step. Firing threshold and rheobase were estimated from voltage responses to 500-ms current ramps ranging from −50 to +200 pA. In voltage-clamp mode, the membrane potential was held at −60 mV; series resistance was compensated (70–80%) and monitored throughout. For VA1d PN recordings that require electrical stimulation of ORN axons, preserving the antennal nerve bundles was critical. To achieve this, the second and third segments of both antennae were carefully removed using forceps before brain dissection, a procedure that usually ensure the complete preservation of the antennal nerve bundles. A suction pipette with an approximately 10 μm opening, filled with extracellular solution, was used to stimulate the antennal nerve by gently pulling the nerve bundle into the pipette. Electrical stimulation was applied by connecting the suction pipette to an isolated pulse stimulator (Model 2100, A-M Systems), which was controlled via a digitizer (Digidata 1550B). The pulse duration was set to 0.1 ms, and the stimulation intensity was increased to the minimum level required to elicit maximal synaptic responses. The external saline was supplemented with GABA receptor antagonists, including 5 μM Picrotoxin and 50 μM CGP 54626 hydrochloride. To assess the spatial coupling between Ca^2+^ channels and synaptic vesicles, we recorded evoked EPSCs from VA1d PNs in response to ORN stimulation at 15-second intervals. After establishing a stable 10-minute baseline, 150 μM EGTA-AM (Sigma-Aldrich), a membrane-permeable slow Ca^2+^ chelator, was bath-applied for 30 minutes. EPSCs were continuously monitored during EGTA-AM treatment. A progressive reduction in EPSC amplitude during EGTA-AM application was interpreted as evidence of loose coupling between Ca²⁺ channels and the synaptic release machinery. For quantification, the amplitude of each EPSC was first measured individually, and the four responses acquired per minute were then averaged to yield a single representative value. To record mEPSCs, tetrodotoxin (TTX, 1 μM) was added to the external saline to block action potentials. The identity of the PN neurons was determined post hoc via immunohistochemistry with a fluorescence conjugated streptavidin (1:1000). For DPM recordings requiring optogenetic activation, 2-ms whole-field light stimulation was delivered through a 40X objective (Olympus LUMPLFLN 40XW) using a ~470 nm CoolLED pE-300ultra light source. The light intensity was fixed at 16 mW/mm², a level that reliably produced saturated synaptic responses. In nicotine uncaging experiments, after achieving whole-cell configuration, 1 μM TTX, 5 μM picrotoxin, 50 μM CGP 54626, and 150 μM PANic2 were bath-applied for 5 minutes to block presynaptic release and isolate postsynaptic responses. A 1-second uncaging pulse was then delivered using a ~405 nm LED, directed through a condenser positioned beneath the recording chamber to uniformly illuminate the brain. For both VA1d PN and DPM recordings, paired-pulse stimuli were applied at defined inter-stimulus intervals (ISIs). Each ISI condition was repeated five times, with a 30-second inter-trial interval to ensure full recovery between stimulations. The data were analyzed in IGOR Pro (v9.0.5.1, WaveMetrics) using a custom-written script. Extracellular Ca2+ concentration for measuring P_r_ is specified in the figures.

For sharp-electrode recordings in the neuromuscular junctions (NMJs), third instar *Drosophila melanogaster* larvae were used, and recordings were made from muscle 6 in abdominal segments A2 to A4 specifically. Dissections were conducted in HL-3 saline composed of (in mM): 110 NaCl, 5 KCl, 10 NaHCO₃, 5 HEPES, 30 sucrose, 5 trehalose, and 10 MgCl₂, adjusted to pH 7.2. Segmental motor nerves were severed during dissection. Recordings were carried out at room temperature (21–23 °C) using sharp intracellular microelectrodes filled with 3 M KCl (resistance <20 MΩ). Only cells with an input resistance ≥3 MΩ and a resting membrane potential more negative than –60 mV were included in the analysis. The membrane potential was held at –70 mV throughout all recordings. Synaptic currents and membrane potentials were recorded using an Axoclamp 900A amplifier and digitized via a Digidata 1440A interface (Molecular Devices). Data were acquired and analyzed using pCLAMP 10 software (Clampfit, Molecular Devices). Excitatory junctional currents (EJCs) were evoked by stimulating the segmental motor nerve at twice the threshold intensity using a suction electrode. EJCs were low-pass filtered at 1 kHz. Paired-pulse stimuli were applied at ISI of 10 and 20 ms, under 1 mM Ca^2+^ concentration.

For whole-cell patch-clamp recordings in mice, acute slices were prepared from 9 weeks old male C57/Bl6 control and sleep deprived (SD) mice. In short, immediately after regular sleep (control) or sleep deprivation (SD) animals were sedated using isoflurane and after decapitation, the brain was quickly removed and transferred into ice-cold cutting solution (in mM): 110 choline chloride, 26 NaHCO_3_, 11.6 Na-ascorbate, 10 D-glucose, 7 MgCl_2_, 3.1 Na-pyruvate, 2.5 KCl, 1.25 NaH_2_PO_4_, 0.5 CaCl_2_; 300–315 mOsm, pH adjusted to 7.4, with 5% CO2/95% O_2_. Slicing was performed on a vibratome (Leica VT1200) and coronal slices of 250 µm were made of the medial prefrontal cortex (mPFC) or barrel cortex (BC). For mPFC slices a cutting angle (~15 degrees) was used to improve neuronal integrity. Immediately after cutting the slices were transferred to 32 °C cutting solution for 6 minutes to recover and afterwards were stored at room temperature in holding solution (in mM): 126 NaCl, 26 NaHCO_3_, 10 D-glucose, 6 MgSO_4_, 3 KCl, 1 CaCl_2_, 1 NaH_2_PO_4_, 295-305mM, pH adjusted to 7.4, with 5% CO_2_/95% O_2_. Slices were stored for ~1 hour before experiments. During experiments brain slices were continuously perfused in a submerged chamber (Warner Instruments) at a rate of 3-4 ml/min with recording solution (in mM) 127 NaCl, 2.5 KCl, 1.25 NaH_2_PO_4_, 25 NaHCO_3_, 1 MgCl_2_, 2 CaCl_2_, 25 glucose (pH 7.4 with 5% CO2/ 95% O2), including bicuculline (20 µM) and DPCPX (1 µM). All recordings were done between 32-33 °C. For mESPC recordings, TTX (1 µM) was added to the recording solution. Whole-cell patch-clamp recordings were done using borosilicate glass recording pipettes (resistance 3.5–5 MΩ, Sutter P-1000) filled with the following internal solution (in mM): 115 CsMSF, 20 CsCl, 10 HEPES, 2.5 MgCl_2_, 4 ATP, 0.4 GTP, 10 Creatine Phosphate and 0.6 EGTA. Alexa594 (15 µM) was added to the internal medium to visually identify pyramidal cells shortly after establishing whole cell configuration. Whole-cell patch-clamp recordings were done using a double EPC-10 amplifier under control of Patchmaster v2 x 32 software (HEKA Elektronik, Lambrecht/Pfalz, Germany). Currents were recorded at 20 kHz and low-pass filtered at 3 kHz when stored. The series resistance was compensated to 70-80% and monitored throughout the experiments. We recorded L2/3 pyramidal cells combined with extracellular stimulation in L1 (mPFC) or in L4 (BC). Spontaneous and miniature input was measured using whole-cell voltage clamp at Vm=-70 mV. For extracellular stimulation (50-120 µA; 0.2 ms (Isoflex, A.M.P. Instruments LTD)), we used theta glass pipettes (borosilicate theta glass, Hilgenberg) filled with recording solution. Paired pulse (10 and 20 ISIs, repeated 10-15 times every 15 seconds) currents were measured in whole-cell voltage clamp at a holding potential of -70 mV. Evoked data were analyzed using Fitmaster (HEKA Elektronik, Lambrecht/Pfalz, Germany), miniature inputs were analyzed using Mini Analysis program (Synaptosoft).

For all recordings in this study, the amplitude of EPSC was determined by measuring the difference between the baseline current immediately preceding the stimulus and the peak of the evoked current. Paired-pulse ratio (PPR) was calculated as the ratio of the second EPSC amplitude to the first.

**Two-photon functional imaging**

Female flies aged 5–8 days post-eclosion were dissected as in the electrophysiological experiments. Isolated brains were then imaged using a two-photon microscope (FemtoSmart Dual microscope, Femtonics). Fluorescence was excited with a Chameleon Ultra II laser tuned to 920 nm. Images were acquired at a pixel resolution of 0.4 × 0.4 μm (Single-bouton imaging) or 0.71 x 0.71 μm (Imaging conducted in VA1d glomerulus) using a 25× water-immersion objective (Nikon Plan Apochromat objective) with 1.1 NA, controlled by MESc v3.5 software (Femtonics). During recordings, 1 μM picrotoxin was included in the external solution to GABAA receptor–mediated inhibitory synaptic transmission. The images were recorded at 10 Hz using a galvanometer-based scanner. Optogenetic stimulation (5-ms single pulse) was delivered six times at 1-minute intervals using a 625 nm LED (Optogenetic module, Femtonics) through the objective. Image processing and analysis were conducted using Fiji software. To correct for potential x–y drift, recorded images were aligned using the Fiji plugin TurboReg (*75*) prior to further analysis. Regions of interest (ROIs) were manually defined. For single-bouton imaging, identification of the presynaptic bouton was guided by a z-stack acquired during the experiment. Only presynaptic boutons that were clearly distinguishable in the x, y, and z dimensions in the z-stack images were included in the analysis. F₀ was defined as the average pixel intensity of the region of interest (ROI) over 40 frames preceding stimulation onset. ΔF was calculated as the difference between the pixel intensity of the ROI in each frame and F₀. To represent the relative change in mean ROI intensity, ΔF was normalized to F_0_, with responses averaged over six stimulations.

**Correlated light and electron microscopy and Tomography**

*Drosophila* brains were dissected in external saline solutions external saline solution identical to that used for electrophysiological recordings and fixed for 1 hour in 4% paraformaldehyde with 0.1% glutaraldehyde in 0.1M phosphate buffer pH 7.4 (PB). Samples were washed three times and immobilized on a round coverslip in a low gelling temperature agarose droplet (3% agarose in PB) to fix the orientation, as described previously (*76*). Imaging and branding were performed as reported(*76*). Briefly, Zeiss LSM980 inverted laser scanning confocal microscope (Carl Zeiss Microscopy GmbH, Germany) equipped with Plan-Apochromat 63×/1.4 NA oil immersion objective was used for imaging. 2D near-infrared branding (NIRB) marks were introduced with a tunable two-photon laser (InSight X3, Spectra-Physics, CA, USA) set at 780nm wavelength at 50% laser power and 2.1 µs dwell time using the bleaching module in ZEN Blue 3.7 (Carl Zeiss Microscopy GmbH, Germany). For higher accuracy in correlating the sample during the approach in our SBF-SEM microscope, innovative branding marks along the Z dimension were added. In detail, after 2D NIRB, a confocal image was acquired at the focal plane targeted for branding. Following visual inspection of the 2D branding marks around the glomerular cells of interest, a Z-stack (10 µm) along this anatomical structure was defined, using a 3.5 x 3.5 µm imaging ROI at the intersection of the 2D branding marks. The imaging parameters for branding along the Z dimension matched those of the conventional 2D NIRB. Nyquist sampling was used along the Z direction. An overview confocal Z-stack of the targeted area was then acquired in confocal mode to control the quality of the 3D NIRB. The imaged samples, which were kept hydrated during the NIRB steps, were post-fixed in 2.5% glutaraldehyde and 4% paraformaldehyde in 0.1M cacodylate buffer at 4°C overnight or until further processing.

Next, the agarose droplets containing the brains were stained first with potassium ferrocyanide-reduced osmium (Electron Microscopy Services, PA, USA) for 1 hour, subsequently with 1% thiocarbohydrazide for 20 minutes and osmicated once more with 2% OsO4 for 30 minutes. This was followed by an overnight incubation in 2% uranyl acetate in water at 4°C. The next day, samples were stained en bloc with lead aspartate and dehydrated in an ascending series of ethanol solutions. Samples were dehydrated completely in pure acetone and subsequently embedded in Agar 100 resin (Laborimpex; Agar Scientific), with the inverted BEEM® Embedding Capsules covering the agarose droplet containing the fixed brain.

The embedded brains were cut out of the resin block and mounted on aluminium pin stubs (Gatan, CA, USA) with conductive CircuitWorks epoxy glue (Chemtronics, GA, USA). Next, a Zeiss Sigma Variable Pressure SBF-SEM (Carl Zeiss Microscopy GmbH, Germany) with 3View technology (Gatan, CA, USA) was used to approach the region of interest. The region of interest was recognized based on the brain morphology and the abovementioned 3D branding marks. The vertical branding marks eased tracking the region even if the sample block was tilted (fig. S9). Once the region of interest was reached, 300 nm consecutive sections were cut using a Leica Enuity ultramicrotome and collected on triple slot grids (Ted Pella Inc, CA, USA). Grids were sputter-coated with carbon (4nm) in a Leica ACE600 (Leica Microsystems GmbH, Germany) and 10nm gold particles were deposit as fiducial markers.

Tomograms were acquired with a JEOL 200kV S/TEM JEM-F200 electron microscope (Jeol, Japan), equipped with a TVIPS TemCam – XF416 cooled CMOS camera (TVIPS, Germany). The tilt series were recorded by SerialEM (*77*) with 1-degree angular increments from -65° to +65° at 25kx magnification (pixel size 0.5nm). Data alignment and reconstruction was performed with IMOD software (*78*).

For the analysis, all synaptic vesicles were manually annotated in 3D using webKnossos(*79*). The center of the T-bar was defined as the point on the presynaptic membrane corresponding to the plane in which the T-bar appeared most structurally complete. Vesicle positions were quantified within a 350 nm radius from the T-bar center (*80*). The Euclidean distance between the centroid of each annotated vesicle and the center of the T-bar was calculated in 3D using a custom MATLAB script. Docked vesicles were defined as those in direct contact with the presynaptic membrane at the T-bar base.

**Statistical analysis**

Statistical analyses were conducted using GraphPad Prism 10. Full details of the statistical tests, including test type, degrees of freedom, and exact *P*-values, are provided in Table S2 and S4.

All statistical comparisons were two-sided. Normality was assessed using the D'Agostino & Pearson test. For comparisons between two groups, Student’s *t*-tests were used for normally distributed data, and Mann–Whitney tests for non-normally distributed data. For multiple-group comparisons, one-way ANOVA with Šidák ’s post hoc test was used for normally distributed data, and Kruskal–Wallis test with Dunn’s post hoc test for non-normally distributed data. Two-way ANOVA with Šidák’s post hoc test was used where appropriate to assess interactions across multiple factors. A linear mixed-effects model was used to assess the effect of S vs SD conditions on vesicle distance from active zone, with animal ID included as a random intercept to account for repeated measurements within each animal.

**Data availability**

The data supporting the findings of this study are available within the article and its Supplementary Materials. Raw datasets generated during the current study are available from the corresponding author upon request. Source data will be provided with publication.

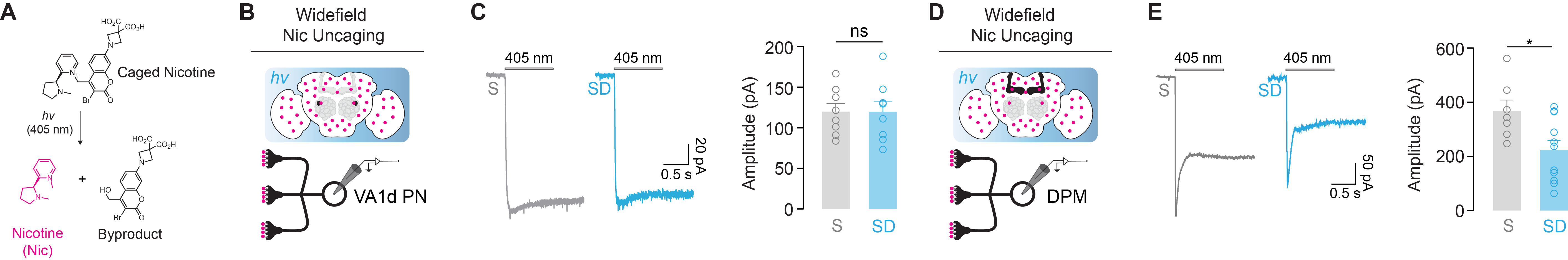

Fig. S1. Circuit-specific cholinergic modulation by sleep deprivation revealed through nicotinic uncaging. (A) Schematic of photolysis of caged nicotine (PANic2), which releases free nicotine and a byproduct upon 405nm illumination. (B) Caged nicotine was bath-applied and uncaged with wide-field 405nm light during recordings from VA1d PNs. (C) Left: Representative photolysis-evoked responses in VA1d PNs from S and SD groups. Right: Photolysis-induced EPSC amplitudes show no significant difference between groups (S: n = 8 neurons; SD: n = 8 neurons). (D) Caged nicotine was bath-applied and photoreleased with widefield 405 nm light during recordings from DPM neurons. (E) Left: Representative photolysis-evoked responses in DPM neurons from S and SD groups. Right: Photolysis-induced EPSC amplitudes are significantly reduced in SD (S: n = 7 neurons; SD: n = 11 neurons). Each data point corresponds to a single-cell recording from an individual fly. Data are presented as mean ± s.e.m. For statistical details, see Table S4. ns, not significant; *P < 0.05, **P < 0.01, ***P < 0.001.

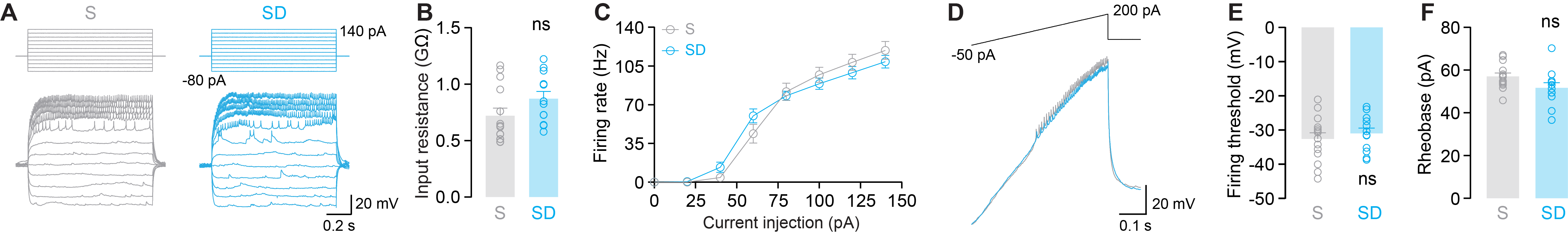

Fig. S2. Intrinsic excitability of Va1d projection neurons is unaffected by sleep deprivation. (A) Representative traces of current-clamp recordings from Va1d projection neurons in flies under sleep (S) and sleep deprivation (SD) conditions in response to step current injections (−80 to +140 pA). (B) Input resistance does not differ significantly between S and SD groups (S: n = 14 neurons; SD: n = 12 neurons). (C) Frequency–current (F–I) relationship showing firing rate as a function of current injection amplitude; no significant difference is observed between groups (S: n = 14 neurons; SD: n = 12 neurons). (D) Example ramp current injection traces to assess spike threshold and rheobase (S: gray trace; SD: blue trace). (E) Firing threshold remains unchanged between S (n=14 neurons) and SD conditions (n=12 neurons). (F) Rheobase current also shows no significant difference between S (n=14 neurons) and SD conditions (n=12 neurons). Each data point corresponds to a single-cell recording from an individual fly. Data are presented as mean ± s.e.m. For statistical details, see Table S4. ns, not significant; *P < 0.05, **P < 0.01, ***P < 0.001.

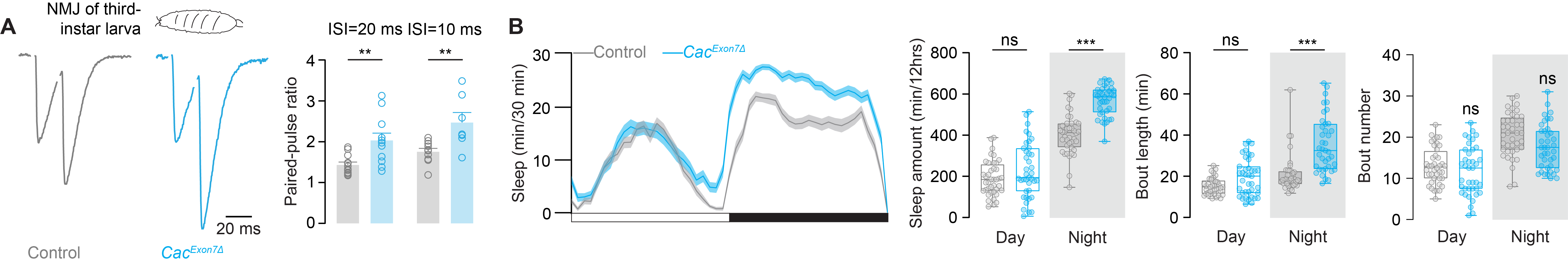

Fig. S3. Cac*^Exon7∆^* reduces presynaptic release probability and promotes nighttime sleep. (A) Paired-pulse recordings at the neuromuscular junction (NMJ) of third-instar larvae. Left: Representative normalized paired EPSCs from Control and *Cac^Exon7∆^* larvae, a hypomorphic VGCC mutant. Right: Paired-pulse ratio is elevated in *Cac^Exon7∆^* larvae at inter-stimulus intervals (ISI) of 20 ms (Control: n = 12 NMJs from 7 flies and *Cac^Exon7∆^* = 11 NMJs from 7 flies) and 10 ms (Control: n = 10 NMJs from 7 flies and Cac^Exon7∆^ = 7 NMJs from 7 flies), under 1.5 mM extracellular Ca^2+^, indicating decreased presynaptic release probability. (B) From left to right: Sleep profiles binned in 30-min intervals; total sleep amount during the day and night; mean sleep bout length during the day and night; and number of sleep bouts during the day and night in control (n = 40 flies) and Cac^Exon7∆^ (n = 39 flies) flies. *Cac^Exon7∆^* flies exhibit significantly increased sleep amount and longer sleep bout length during the night compared to controls. Data presented in panel A is mean ± s.e.m. Box plots in panel B display the median (center line), 25th and 75th percentiles (box), and minimum to maximum values (whiskers); Data point represents a single fly as indicated. For statistical details, see Table S4. ns, not significant; *P < 0.05, **P < 0.01, ***P < 0.001.

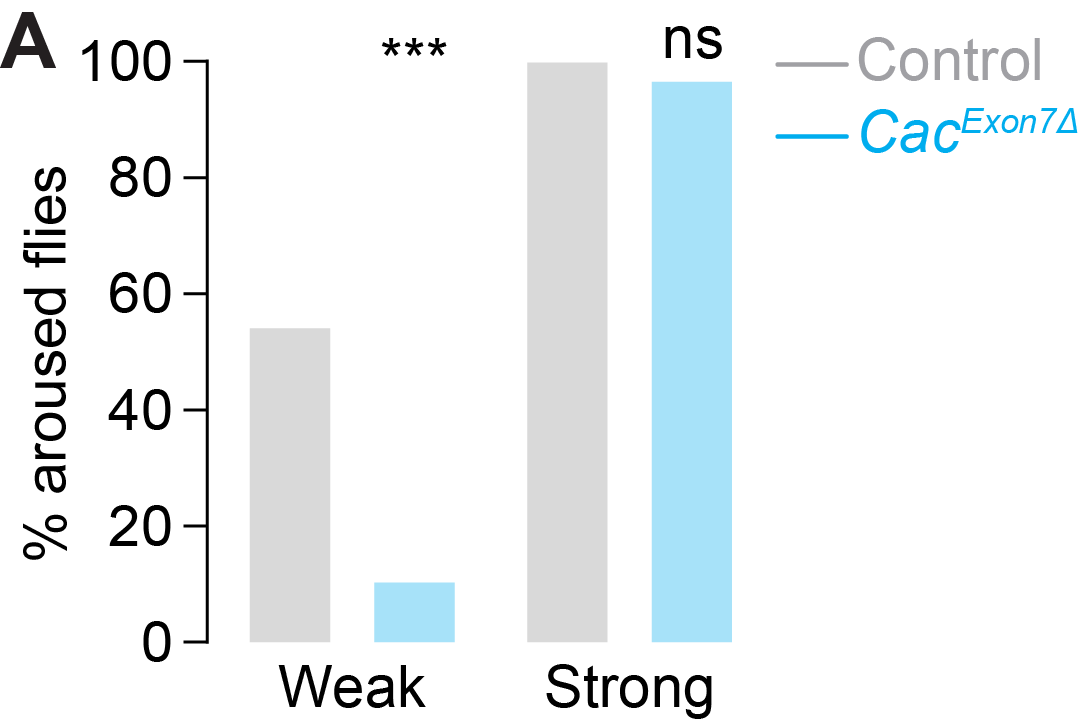

Fig. S4. *Cac^Exon7Δ^* exhibited increases arousal threshold. (A) Percentage of flies that were aroused from sleep in response to weak or strong mechanical stimulation. Flies carrying the *Cac^Exon7∆^* mutation (blue) show a significantly reduced arousal response to weak stimuli compared to controls (grey), while responses to strong stimuli remain unaffected. Different groups of flies were tested for weak (Control: n=46 flies ; *Cac^Exon7∆^*: n=57 flies) and strong stimuli (Control: n=26 flies ; *Cac^Exon7∆^* n=31 flies). For statistical details, see Table S4. ns, not significant; *P < 0.05, **P < 0.01, ***P < 0.001.

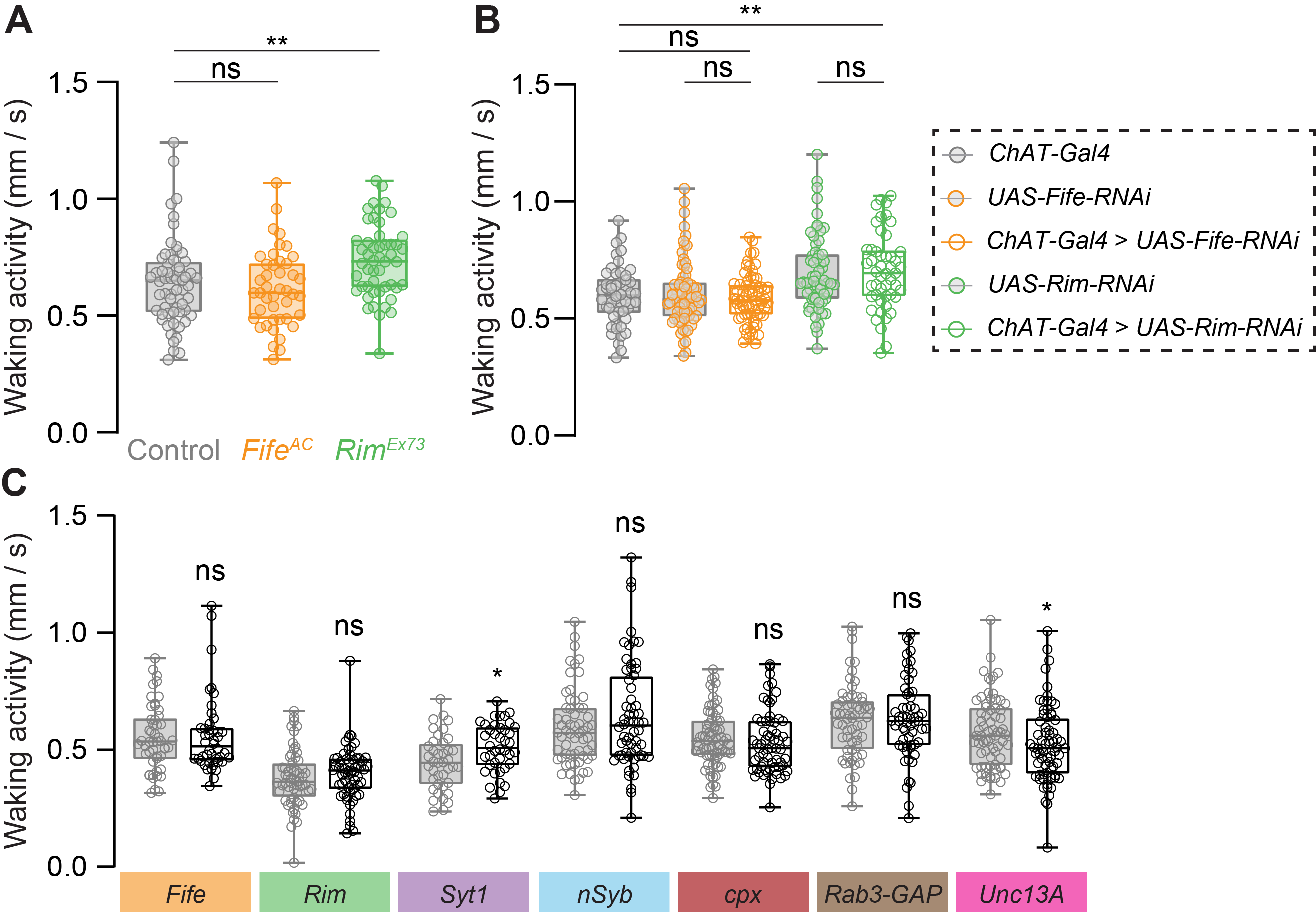

**Fig. S5. Disruption of presynaptic genes minimally affects waking locomotor activity.** (**A**) Waking activity in mutant flies lacking functional Fife (*Fife^AC^*) or Rim (*Rim^Ex73^*), compared to genetic controls. Data are shown for *Fife^AC^* (n=43 flies), *Rim^Ex73^* (n=53 flies) and control (n=60 flies). (**B**) Waking activity in flies with RNAi-mediated knockdown of Fife or Rim specifically in cholinergic neurons using the ChAT-Gal4 driver, along with relevant parental controls. Genotypes shown: *ChAT-T2A-Gal4/+* (n = 50), *UAS-Fife-RNAi/+* (n = 70 flies), *ChAT-T2A-Gal4 > UAS-Fife-RNAi* (n = 68 flies),  *UAS-Rim-RNAi/+* (n = 49 flies), and *ChAT-T2A-Gal4 > UAS-Rim-RNAi* (n = 60 flies). (**C**) Waking activity following adult-specific, pan-neuronal RNAi knockdown of presynaptic genes using the *ELAV-GS* driver with RU486 induction. Each colored box indicates a different gene targeted: *Syt1* (Vehicle, n = 38 flies; RU486, n = 42 flies), *nSyb* (n = 60, 57 flies), *cpx* (n = 75, 64 flies), *Rab3-GAP* (n = 60, 59 flies), *Unc13A* (n = 69, 68 flies), *Fife* (n = 50, 44 flies), *Rim* (n = 59, 60 flies), and *dlg1* (n = 28, 30 flies). For statistical details, see Table S4. *P < 0.05, **P < 0.01, ***P < 0.001.

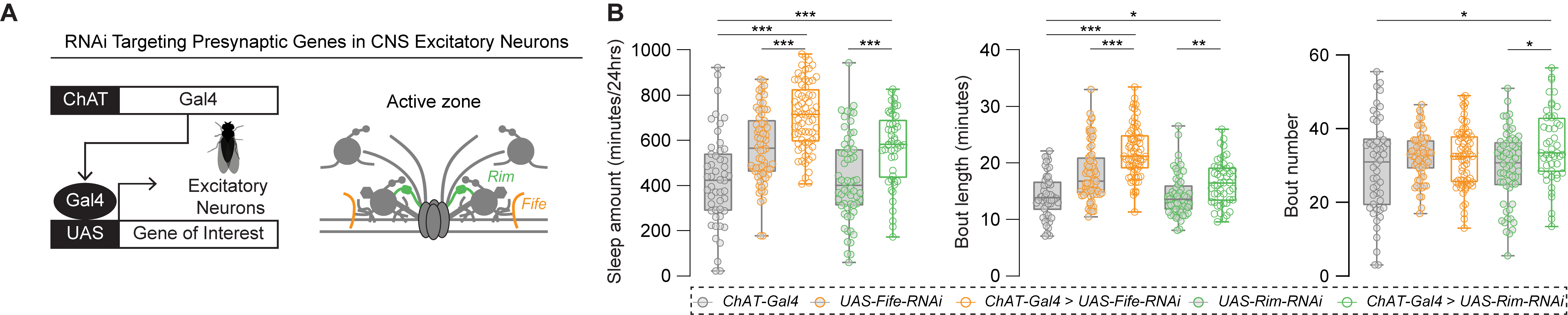

**Fig. S6. Targeted knockdown of Rim and fife in cholinergic neurons enhances sleep and alters sleep architecture.** (**A**) Experimental strategy for targeting gene knockdown to central excitatory neurons using *ChAT-T2A-Gal4* driving *UAS-RNAi* expression. Right: Schematic highlighting the presynaptic active zone with Rim and Fife indicated. (**B**) Sleep architecture. Left to right: Total sleep duration (min/24 hrs), sleep bout length (min), and number of sleep bouts. Genotypes shown: *ChAT-T2A-Gal4/+* (n = 50), *UAS-Fife-RNAi/+* (n = 70 flies), *ChAT-T2A-Gal4 > UAS-Fife-RNAi* (n = 68 flies),  *UAS-Rim-RNAi/+* (n = 49 flies), and *ChAT-T2A-Gal4 > UAS-Rim-RNAi* (n = 60 flies). Box plots display the median (center line), 25th and 75th percentiles (box), and minimum to maximum values (whiskers); Each data point represents a single fly as indicated. For statistical details, see Table S4. *P < 0.05, **P < 0.01, ***P < 0.001.

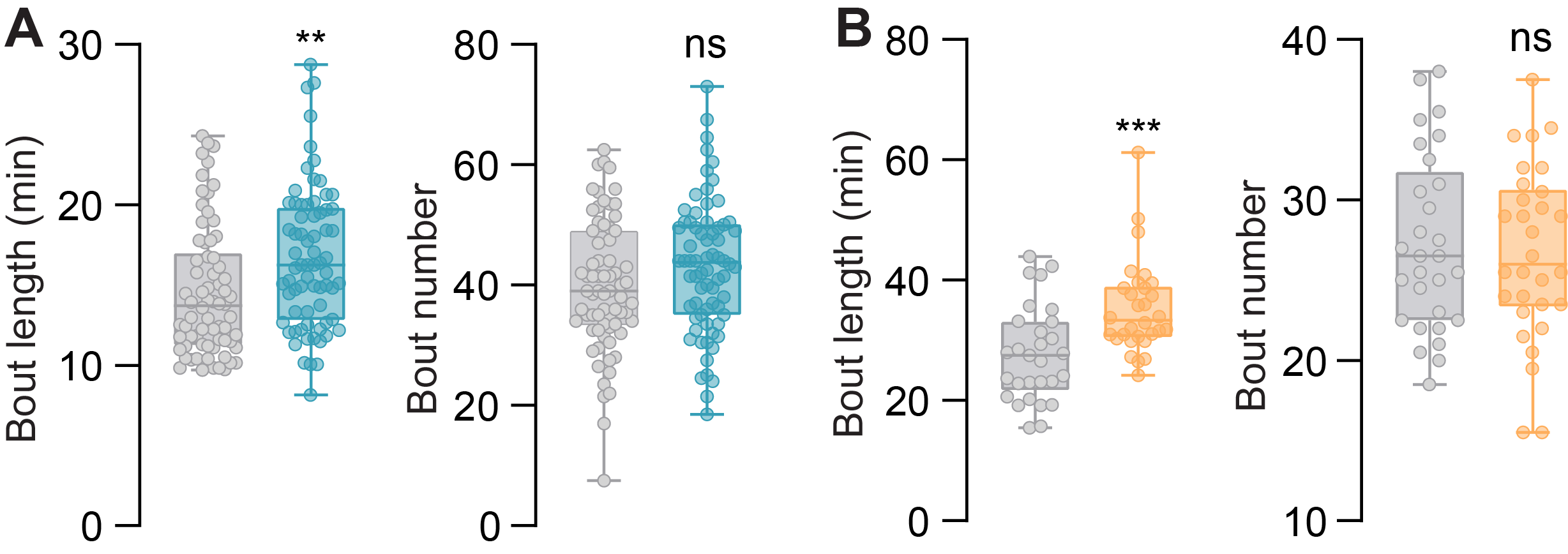

**Fig. S7. Sleep architecture changes following pan-neuronal induction of *Sik3^S3583A^* and *PKAR* in adult flies.** (A) Sleep architecture in flies expressing *Sik3^S3583A^* under *ELAV-GS* control. RU486-fed flies show significantly increased sleep bout length compared to vehicle controls, with no significant change in bout number. (Vehicle: n = 70 flies; RU486: n = 70 flies). (B) Sleep architecture in flies expressing *PKAR* under *ELAV-GS* control. RU486-fed flies (orange) also exhibit significantly increased bout length, while bout number remains unchanged (Vehicle: n=29 flies; RU486: n=30 flies.). Box plots in panels A and B display the median (center line), 25th and 75th percentiles (box), and minimum to maximum values (whiskers); Each data point represents a single fly as indicated. For statistical details, see Table S4. *P < 0.05, **P < 0.01, ***P < 0.001.

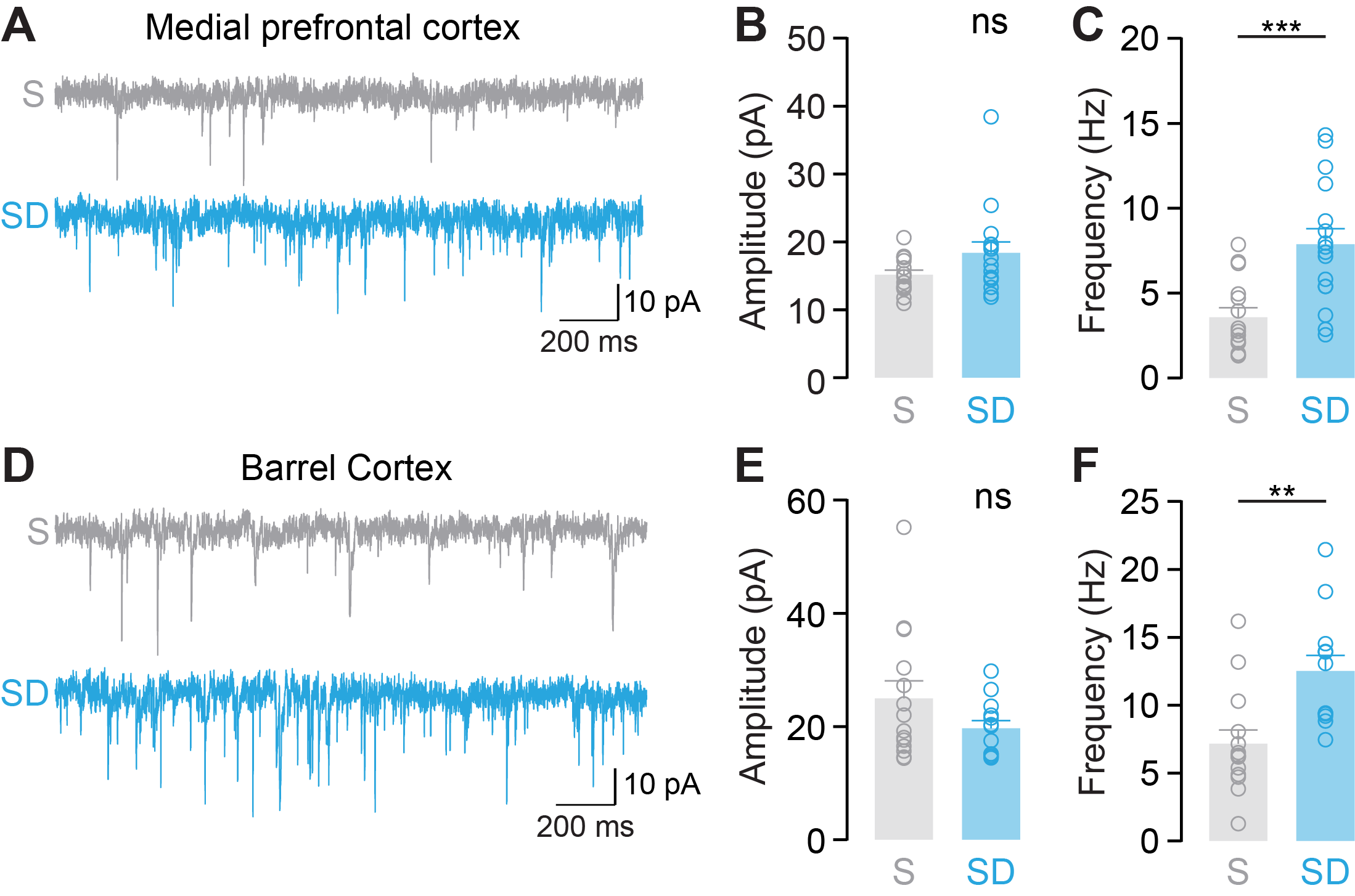

**Fig. S8. Sleep deprivation increases the frequency of miniature excitatory synaptic events in mouse cortical pyramidal neurons.** (**A**) Representative mEPSC traces from layer2/3 pyramidal neurons in the anterior cingulate cortex (ACC) of sleep (S) and sleep-deprived (SD) mice. (**B**) Mean mEPSC amplitude (S: n = 15 neuron from 5 mice; SD: n = 16 neurons from 5 mice). (**C**) Mean mEPSC frequency is significantly increased in SD (n = 15 neurons from 5 mice) compared to S (n =16 neuron from 5 mice). (**D**) Representative mEPSC traces from pyramidal neurons in the layer 4 barrel cortex of S and SD mice. (**E**) Mean mEPSC amplitude (S: n = 13 neuron from 4 mice; SD: n = 14 neurons from 5 mice). (**F**) Mean mEPSC frequency is significantly increased in SD (n = 14 neurons from 5 mice) compared to S (n = 13 neuron from 4 mice). For statistical details see Table S4. ns, not significant; *P < 0.05, **P < 0.01, ***P < 0.001.

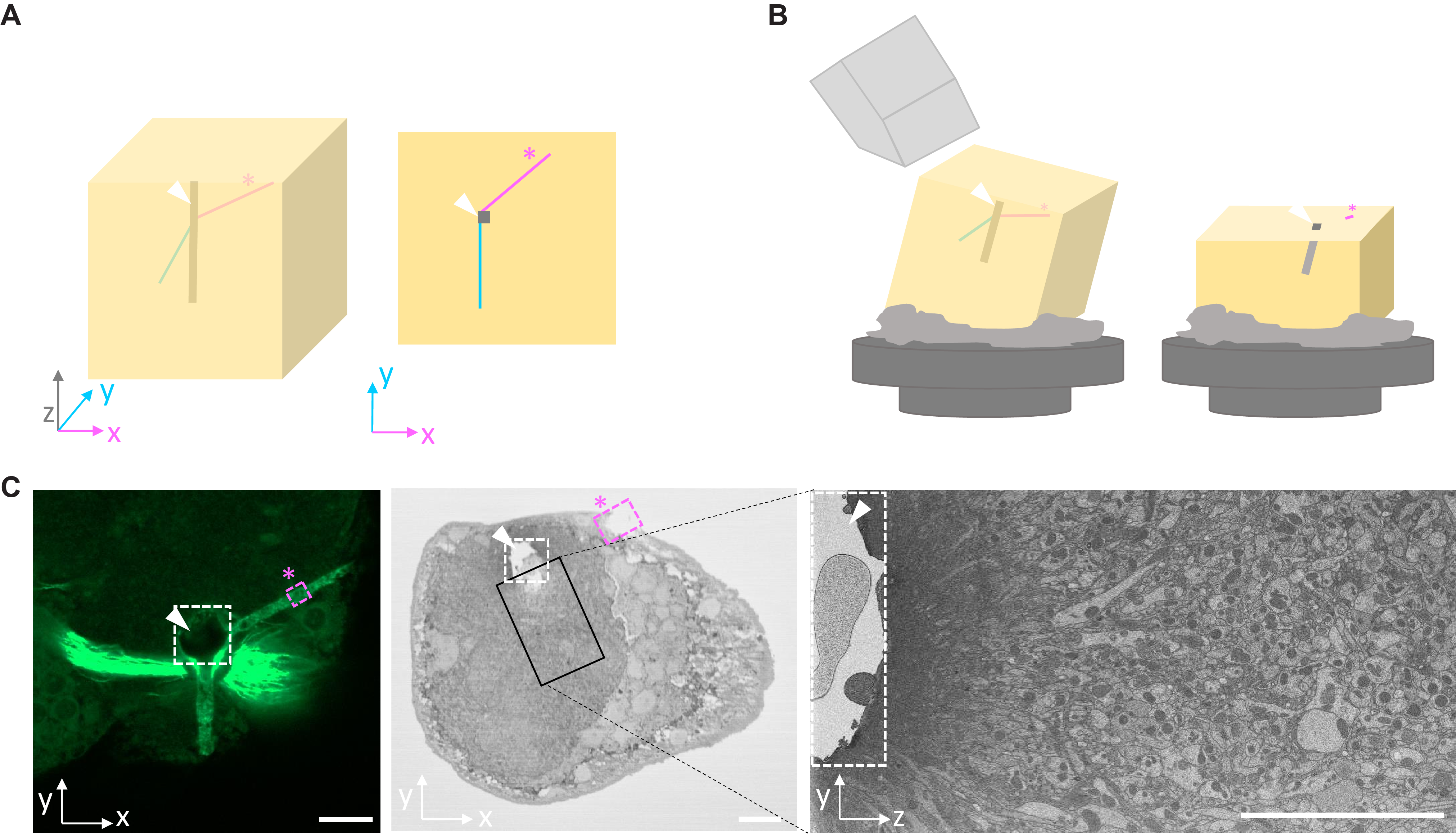

**Fig. S9. 3D NIRB facilitates tracking region of interest (ROI) during approaching in serial block face-SEM (SBF-SEM**). (**A**) Schematic representation of front and top view of a resin block with 2D and 3D branding marks, with the X- and Y-axis colored in magenta and blue respectively, and the Z-axis represented as a thicker grey line. (**B**) Schematic representation of approaching with SBF-SEM. Even if the block is tilted on a pin, the vertical branding mark will remain visible across the trimmed volume. (**C**) Fluorescence (left) and backscatter electron microscopy images of 2D and 3D branding marks next to glomerulus of interest (XY projection in center, YZ projection in right panel). The white arrow head indicates the vertical branding mark, while the asterisk highlights the location of the conventional 2D marks. Scale bar 10µm.

**Table S1. Genotypes and experimental conditions corresponding to Fig. 1–4.**

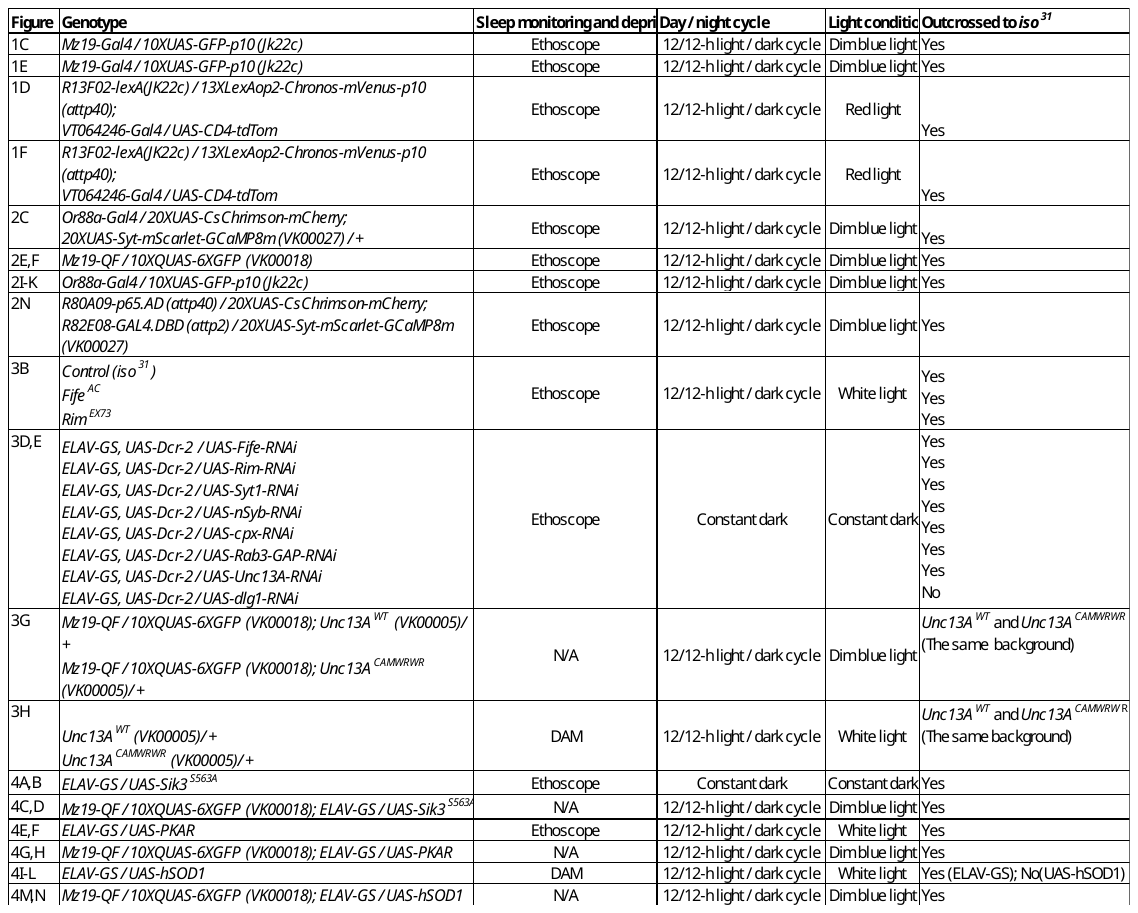

**Table S2. Statistical summary corresponding to Fig. 1–4.**

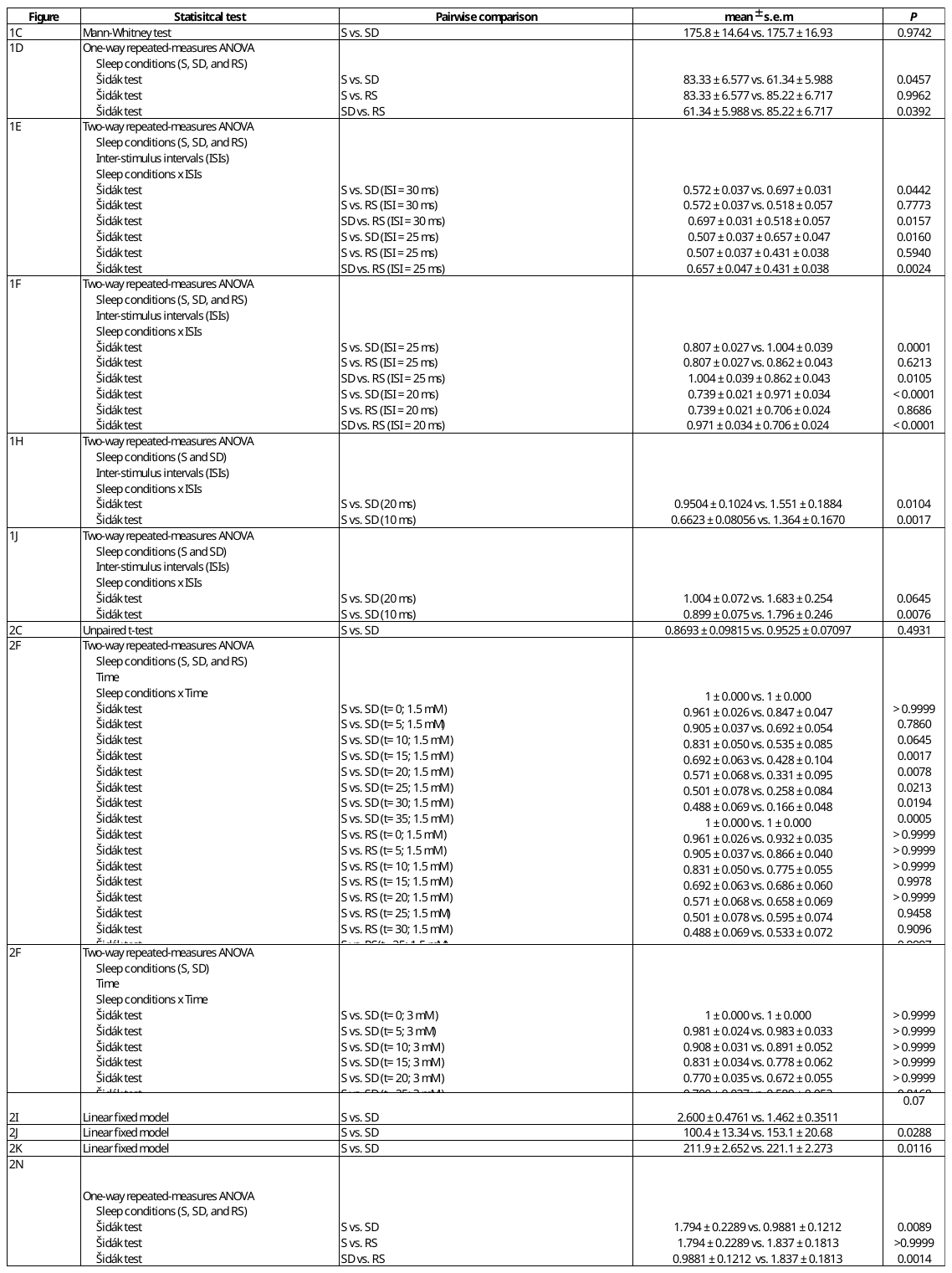

**Table S3. Genotypes and experimental conditions corresponding to Fig. S1-S7.**

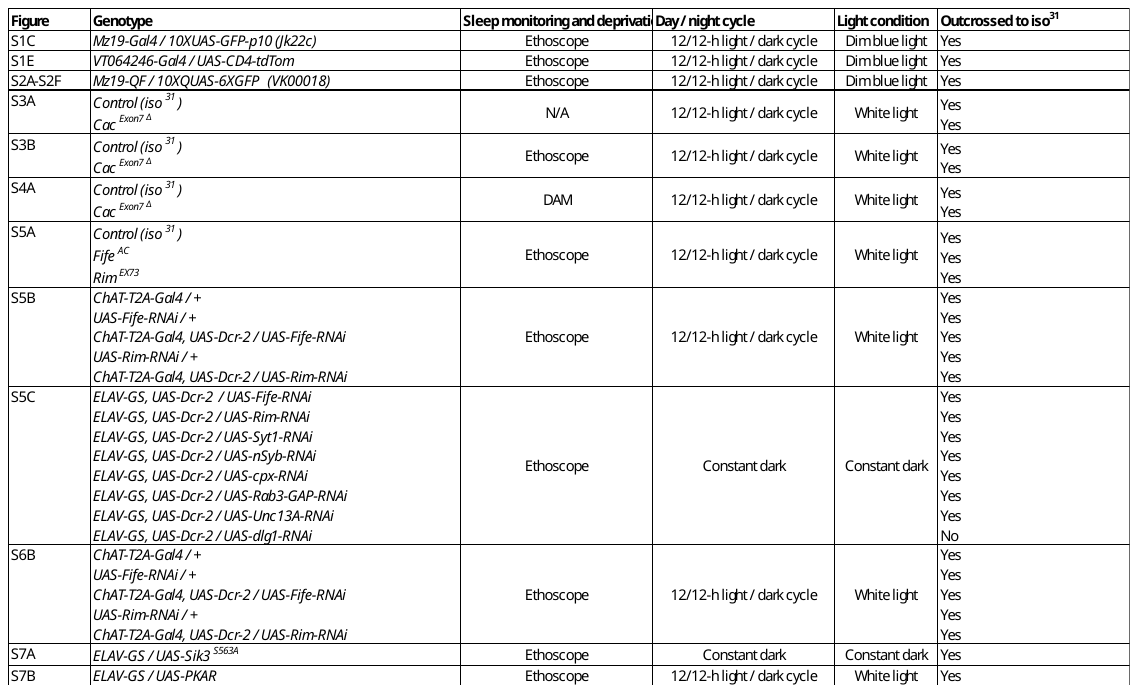

**Table S4. Statistical summary corresponding to Fig. S1–S8.**

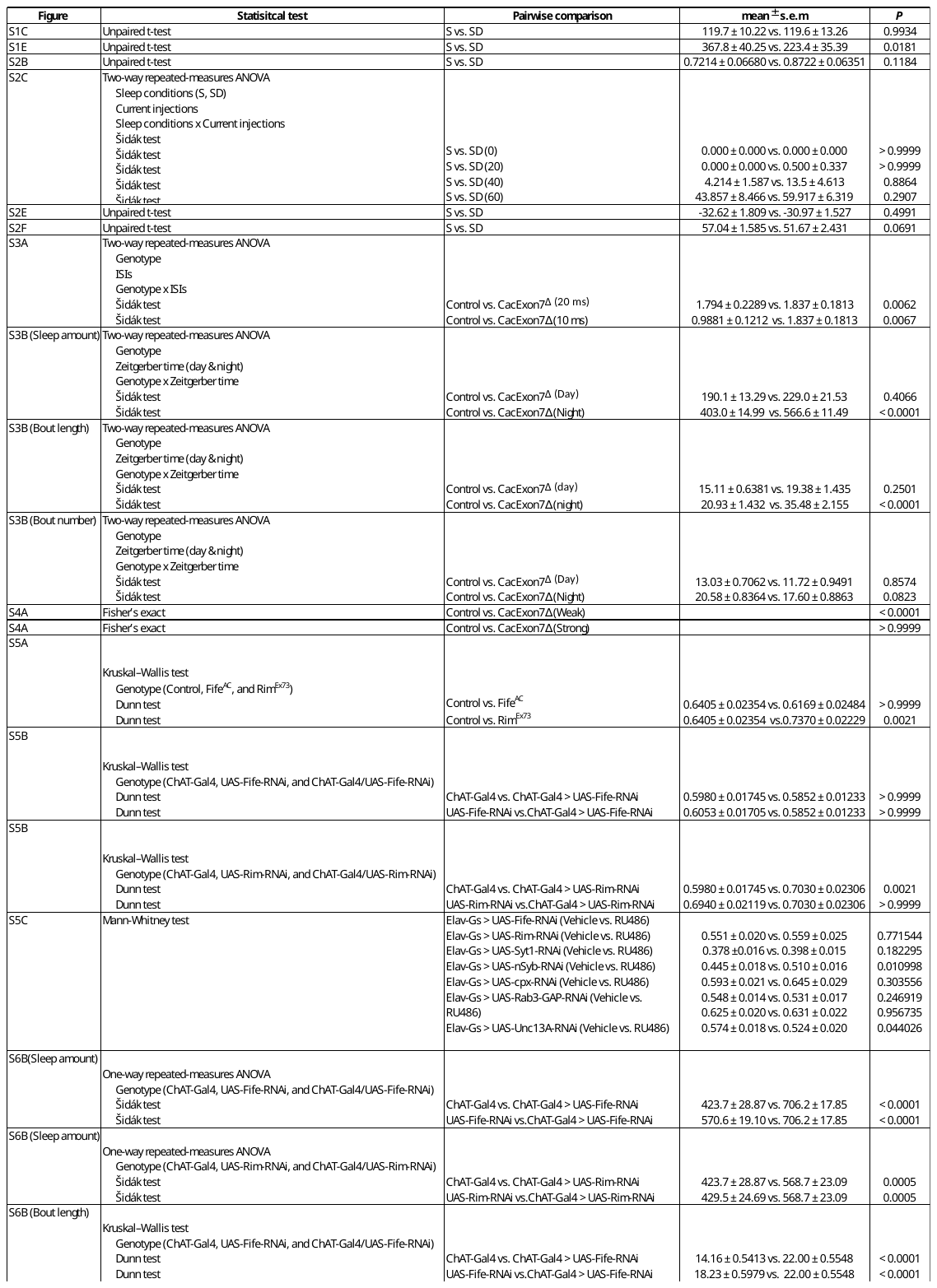

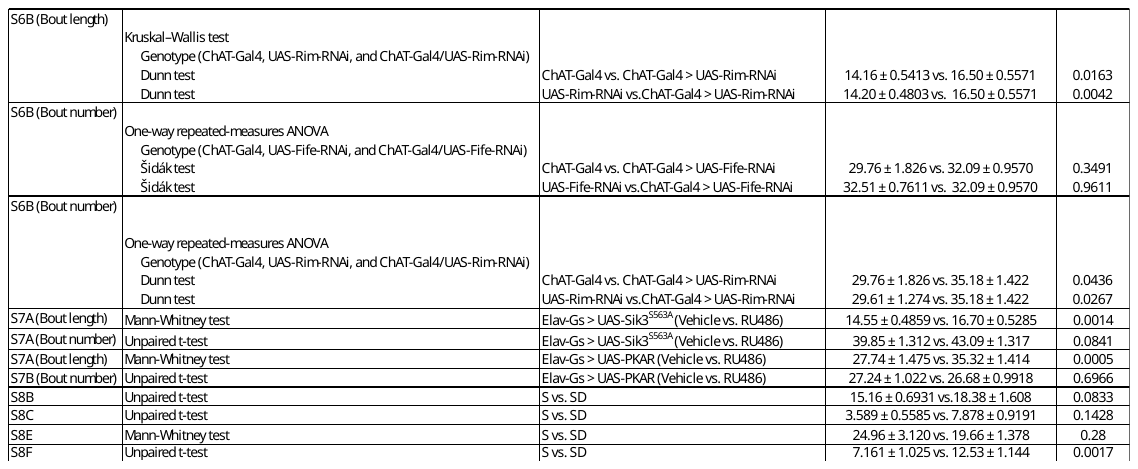
